# PACAP engages distinct signaling programs across cortical neuronal and non-neuronal cell types

**DOI:** 10.64898/2026.10.25.761643

**Authors:** Salome Nora Heimbach, Álvaro Collazos Matute, Bayla Dolman, Victoria Bald, Sarah Melzer

**Affiliations:** Medical University of Vienna, Center for Brain Research, Division of Neuronal Cell Biology

## Abstract

Neuropeptides form modulatory communication channels that tune neuronal circuits and their functions, often through selective targeting of distinct cell populations. Pituitary adenylate cyclase-activating polypeptide (PACAP) is an important neuropeptidergic regulator of fear and stress. However, its receptors are surprisingly ubiquitous across most brain areas and cell types, making the physiological roles and functional mechanisms difficult to predict. Here, we investigate the cell type-specific effects of PACAP in the temporal association cortex (TeA), a brain region implicated in fear learning and memory. *Adcyap1r1*, encoding the PACAP receptor PAC1, was broadly expressed across the majority of neuronal and non-neuronal cells. Yet, PACAP engaged highly diverse molecular pathways across functionally distinct cell types that were differentially recruited during auditory fear learning and retrieval, providing a framework for how broad receptor expression can yield specific network modulation. Our data underscore the importance of dissecting cellularly resolved peptidergic signaling to understand functional mechanisms of neuropeptides.

## Main

Neuromodulatory systems play central roles in many brain functions, often by modulating circuit activity according to affective and cognitive states such as arousal, stress, and attention. Pituitary adenylate cyclase-activating polypeptide (PACAP) and its high-affinity receptor PAC1, encoded by *Adcyap1r1*, are widely distributed across the brain (Hannibal, 2002; Hirabayashi et al., 2018; Shioda et al., 1997) and have been implicated in synaptic plasticity, stress adaptation and fear-related behavior (Hammack et al., 2009; Hirabayashi et al., 2018; Otto et al., 2001; Rajbhandari et al., 2023; Ressler et al., 2011; Schmidt et al., 2015). In fear-circuits, including the hippocampus, amygdala and prefrontal cortex, PACAP-PAC1 signaling can modulate activity-dependent gene expression, synaptic efficacy and memory consolidation (Hirabayashi et al., 2018; Kirry et al., 2018; Meloni et al., 2016; Rajbhandari et al., 2023; Schmidt et al., 2015). PACAP signaling has been linked to stress disorders and sex-dependent vulnerability in human studies associating *ADCYAP1R1* variants and circulating PACAP levels with increased risk for post-traumatic stress disorder (PTSD) symptoms in females (King et al., 2017; Mercer et al., 2016; Ressler et al., 2011). These studies identify PACAP-PAC1 signaling as a broadly acting and functionally important modulatory pathway.

Although several regions have been implicated in the role of PACAP-PAC1 signaling in fear and stress responses (Kirry et al., 2018; Meloni et al., 2016; Schmidt et al., 2015), the precise cellular and circuit-level mechanisms linking receptor activation to behavioral outcomes remain largely elusive. In contrast, a growing number of *in vitro* studies suggests high diversity of PAC1-signaling pathways. Thus, PAC1 activation in Xenopus oocytes, cultured cortical neurons and astrocytes, and in somatosensory cortex pyramidal cells can engage cAMP production (Grimaldi & Cavallaro, 1999; Spengler et al., 1993), a common downstream effect of Gs-coupled receptors and adenylyl cyclase activation. However, in mouse embryonic fibroblast cell culture and cortical neuronal cell culture, PAC1 can also evoke phospholipase C (PLC) recruitment and inositol 1,4,5-trisphosphate (IP_3_) and Ca^2+^ mobilization (Grimaldi & Cavallaro, 1999; Pisegna & Wank, 1996), which are most commonly recruited by Gq-coupled receptors. In addition, PAC1 can recruit β-arrestin-dependent receptor internalization and extracellular signal-regulated kinase (ERK) signaling in HEK293T cell culture and dentate gyrus cells (Johnson et al., 2020; Shintani et al., 2018). More recent work identified PAC1-dependent recruitment of voltage-dependent Ca^2+^ channels through unknown mechanisms in prelimbic cortical astrocytes (Cui et al., 2025). Whether this diversity is primarily driven by differences in experimental conditions and the brain areas investigated, or whether cells within the same brain area and under the same physiological conditions respond to PACAP through differential recruitment of intracellular pathways, remains unknown. Resolving this question could provide important insights into how a ubiquitously expressed receptor achieves sufficient specificity to modulate neuronal networks, even when acting on functionally diverse cell types.

Fear learning requires assigning negative value to initially neutral sensory cues. The temporal association area (TeA) is well positioned for this function. It receives convergent sensory and affective input from auditory cortex, thalamus and amygdala, contains multisensory-responsive neurons, and projects to limbic and striatal target areas implicated in threat learning and behavioral expression (Cheng et al., 2025; Dahl et al., 2009; Dalmay et al., 2019; Luo et al., 2022; Zingg et al., 2014). Consistent with this anatomical organization, temporal cortex activity and output pathways are required for auditory threat memory formation and retrieval, and temporal cortex neurons undergo learning-dependent transcriptional and representational changes after fear conditioning (Cho et al., 2016; Dalmay et al., 2019; Luo et al., 2022). Yet, the role and underlying mechanisms of PACAP-PAC1 signaling in the TeA remain unexplored, despite the evolutionarily conserved role of this signaling pathway in fear-related processing (Ressler et al., 2011).

Here we asked how PACAP-PAC1 signaling is organized across neuronal and non-neuronal populations in the TeA, and how PAC1-expressing neurons are engaged during auditory fear learning and memory. We combined cell-type-specific expression mapping, genetically encoded Ca^2+^ and cAMP sensors, new tool development for astrocytic cAMP imaging, pharmacological pathway dissection, circuit tracing, electrophysiology, behavioral testing and *in vivo* optical recordings. This approach allowed us to define PACAP-responsive cortical cell types, identify cell-type-specific intracellular signaling routes, and determine fear learning and memory-associated activity patterns of PAC1-expressing TeA neurons. Together, our findings reveal a cell-type-specific PACAP signaling architecture in the TeA that provides a framework for understanding how neuropeptide signaling may modulate neuronal and astrocytic signal processing during emotional learning and memory.

## Results

### *Adcyap1r1* is expressed across multiple neuronal and non-neuronal cell types in cortex

Previous anatomical studies demonstrated *Adcyap1r1* expression in both glutamatergic and GABAergic neurons in multiple cortical regions (Zhang et al., 2021), and emerging evidence suggests cortical expression in astrocytes (Cui et al., 2025; Heimbach et al., 2026). However, the relative prevalence of *Adcyap1r1* across major molecularly defined cortical inhibitory neuron subclasses, excitatory neurons, and astrocytes has not been systematically quantified. We therefore used fluorescence in situ hybridization (FISH) to assess *Adcyap1r1* co-expression with markers of *Pvalb*^+^, *Sst*^+^, and *Vip*^+^ inhibitory neurons, *Slc17a7*^+^ excitatory neurons, and *Aqp4*^+^ astrocytes in the TeA and additional cortical regions (Fig. 1A-D). *Adcyap1r1* was detected in a high proportion of all three inhibitory neuron subclasses and in 95.2 ± 1.7% of *Aqp4*^+^ astrocytes in the TeA (Fig. 1A-C). In contrast, expression in *Slc17a7*^+^ excitatory neurons was more variable across regions, ranging from ∼42% in prelimbic cortex (PL) to ∼86% in the TeA (Fig. 1B-C). Since excitatory neurons constituted 88.67% among the counted neurons in TeA, the absolute number of *Adcyap1r1*-expressing *Slc17a7*^+^ neurons was higher than for other neuronal cell types (Fig. 1D).

**Fig. 1:**
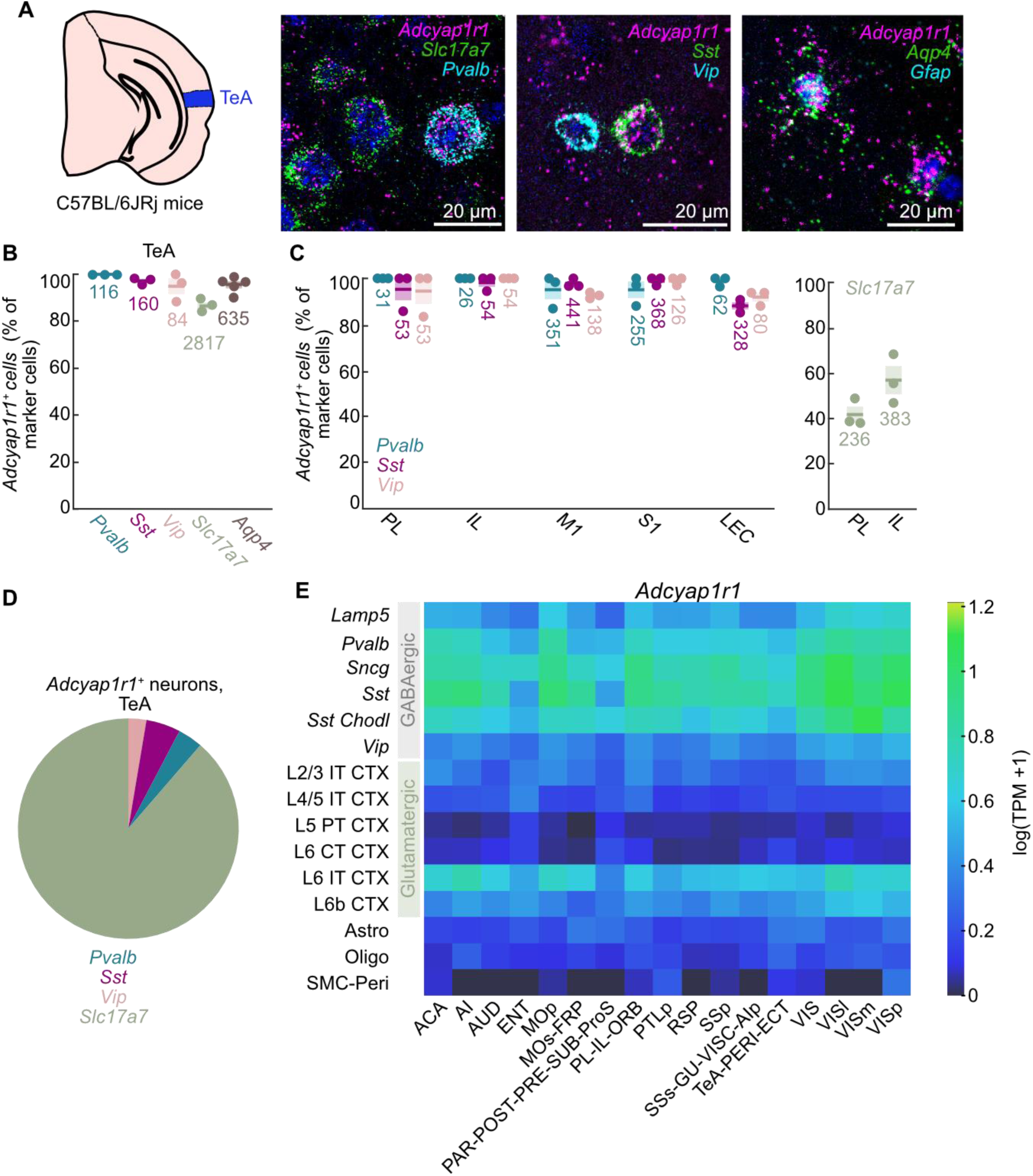
*Adcyap1r1* is expressed across multiple neuronal and non-neuronal cell types in cortex. **A.** Schematic of TeA and representative fluorescence in situ hybridization (FISH) images labeled for *Adcyap1r1* together with cell-type markers (*Slc17a7*, *Sst*, *Pvalb, Vip, Aqp4, Gfap*) and nuclei (DAPI). **B.** Quantification of proportions of indicated cell types expressing *Adcyap1r1* in the TeA. Each dot represents one mouse, absolute number of total analyzed *Adcyap1r1*^+^ cells indicated. *N* = 4 slices per mouse, 3-5 mice. **C.** Quantification of *Adcyap1r1* co-expression with indicated inhibitory neuron (left panel) and excitatory neuron (right panel) marker genes in the indicated brain areas. Absolute number of cells indicated. *N* = 4 slices per mouse, 3 mice. **D.** Percentage of *Adcyap1r1^+^*cells in the TeA co-expressing the indicated inhibitory or excitatory neuron markers (*Adcyap1r1^+^Pvalb^+^* 3,65%; *Adcyap1r1^+^Sst^+^* 5,04%; *Adcyap1r1^+^Vip^+^* 2,64%; *Adcyap1r1^+^Slc17a7^+^* 88,67%). **E.** Heatmap showing expression (log(TPM+1)) of *Adcyap1r1* across major GABAergic, glutamatergic and non-neuronal cell types in 18 cortical area groups. Expression data and cell-type classification were extracted and reanalyzed from Yao et al 2021. **Abbreviations**: ACA = anterior cingulate area; AI = agranular insular area; AIp = agranular insular area, posterior part; AUD = auditory areas; ECT = ectorhinal area; ENT = entorhinal area; GU = gustatory areas; IL = infralimbic cortex; LEC = lateral entorhinal cortex; M1 = primary motor cortex; MOp = primary motor area; MOs = secondary motor area; FRP = frontal pole, cerebral cortex; ORB = orbital area; PAR = parasubiculum; PL = prelimbic cortex; POST = postsubiculum; PRE = presubiculum; ProS = prosubiculum; PTLp = posterior parietal association areas; RSP = retrosplenial area; S1 = primary somatosensory cortex; SSp = primary somatosensory area; SSs = supplemental somatosensory area; SUB = subiculum; TeA = temporal association area; PERI = perirhinal area; VISC = visceral area; VIS = visual areas; VISl = lateral visual area; VISli = laterointermediate area; VISm = medial visual area; VISp = primary visual area. Data are shown as mean ± SEM.

To validate our findings and systematically assess *Adcyap1r1* expression across cortical cell classes, we reanalyzed single-cell RNA sequencing data from Yao et al. (2021) encompassing multiple cortical regions. This analysis confirmed widespread *Adcyap1r1* expression across major GABAergic, glutamatergic and non-neuronal cell populations, with particularly prominent expression in inhibitory neuronal subclasses (Fig. 1E).

In contrast, expression of the PACAP-encoding gene *Adcyap1* was more restricted and enriched within layer 5 glutamatergic neurons (Extended Data Fig. 1A). Together, these findings suggest that PACAP release may originate from defined excitatory neuronal subsets, whereas PAC1 receptor expression is broadly distributed across cortical circuits, positioning PACAP signaling as a widespread modulator of neuronal and astrocytic function.

### Cortical *Adcyap1r1*-Cre^+^ TeA neurons are modulated by PACAP and encode fear memory

To determine whether *Adcyap1r1*^+^ neurons in the TeA express functional levels of PACAP receptors, we first examined intracellular signaling in acute brain slices from Adcyap1r1-Cre mice. Previous cell culture studies suggest that PACAP receptors can couple to both Gs-mediated cAMP signaling, and Gq-mediated Ca^2+^ signaling (Grimaldi & Cavallaro, 1999; Spengler et al., 1993). We first employed the fluorescent cAMP sensor cyclic AMP Difference Detector in situ (cADDis), using viral delivery of AAV-hSyn-DIO-cADDis to TeA *Adcyap1r1*-Cre^+^ neurons (Fig. 2A). Epifluorescent imaging of acute brain slices revealed fluorescence increases to bath application of 300 nM PACAP-38 (hereafter referred to as PACAP) in comparison to vehicle application (*p* = 0.041). Subsequent forskolin application confirmed the viability of the imaged neurons 10 min after PACAP application (Fig. 2B). Importantly, 150 nM of the selective PAC1 agonist maxadilan evoked comparable cAMP increases (Extended Data Fig. 2A*, p* = 0.356), suggesting that the observed transients were predominantly mediated by PAC1 activation. By contrast, when AAV-CBA-DIO-GCaMP6s was expressed in TeA *Adcyap1r1*-Cre^+^ neurons to image Ca^2+^ dynamics, PACAP did not evoke significant fluorescence increases (Fig. 2C-D). Thus, under these conditions, acute PACAP receptor activation in TeA *Adcyap1r1*-Cre^+^ neurons was associated with robust cAMP signaling but not measurable Ca^2+^ transients, suggesting Gs-but not Gq protein coupling.

**Fig. 2:**
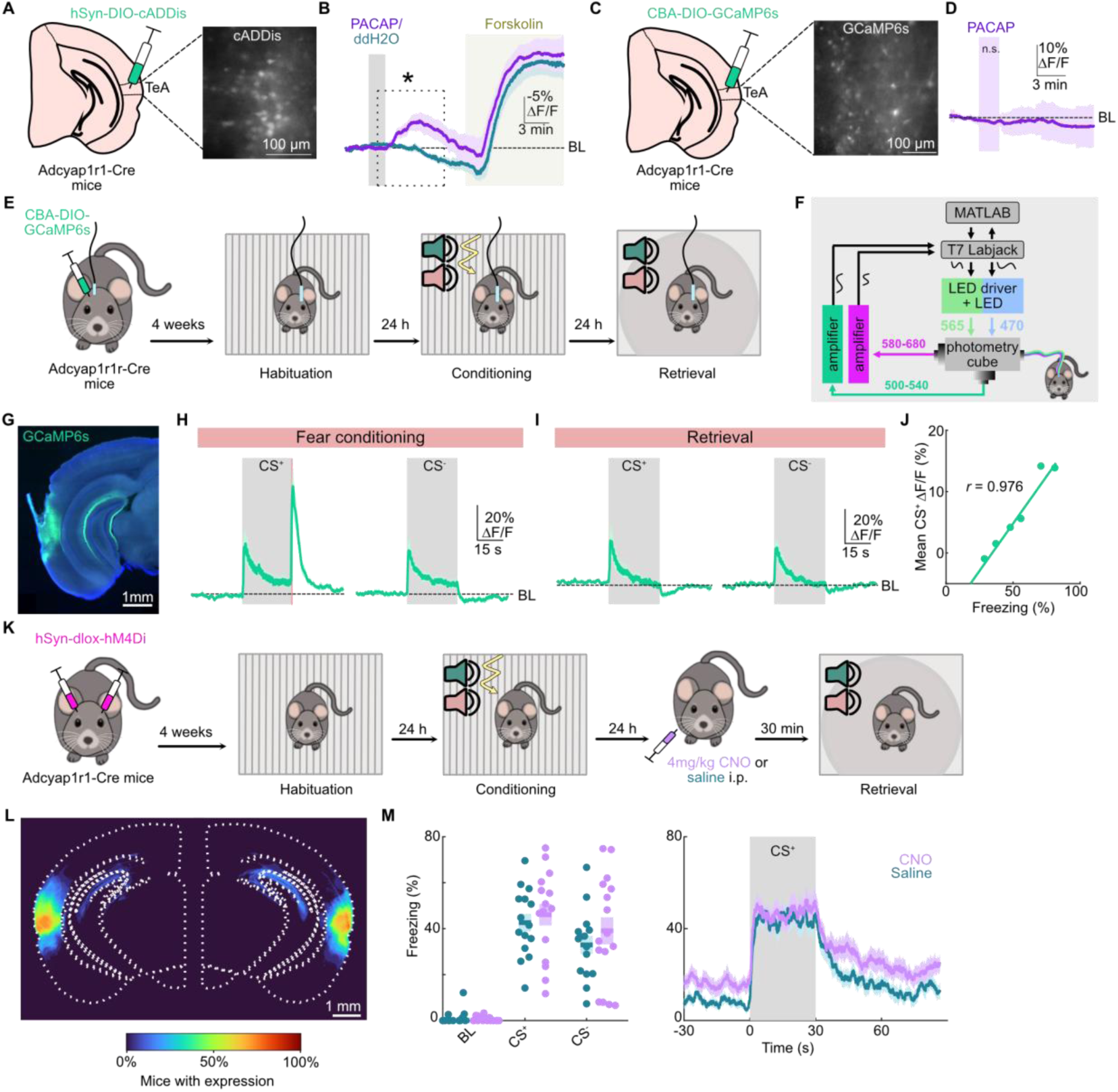
*Adcyap1r1*-Cre^+^ TeA neurons show PACAP-evoked cAMP signaling and are recruited during auditory fear learning and retrieval. **A.** Schematic of AAV-injections of AAV-hSyn-DIO-cADDis into TeA in Adcyap1r1-Cre mice. Exemplary epifluorescence image of cADDis labeling in an acute TeA slice. **B.** Averaged fluorescence cADDis signals (-ΔF/F) of TeA *Adcyap1r1*-Cre^+^ neurons upon PACAP (300 nM) and forskolin (10 µM) application. Peak signals: PACAP -34.11 ± 6.7% ΔF/F; ddH2O control: -13.27 ± 2.5% ΔF/F, *t*(7.68) = 2.912, *p* = 0.041; Welch’s unpaired *t*-test, *N* = 7-8 slices per condition. **C.** Schematic of AAV-injections of AAV-CBA-DIO-GCaMP6s into TeA in Adcyap1r1-Cre mice. Exemplary epifluorescence image of GCaMP6s labeling in an acute TeA slice. **D.** Averaged fluorescence Ca^2+^ signals (ΔF/F) of *Adcyap1r1*-Cre^+^ neurons to PACAP application. Peak signals to PACAP: 1.16 ± 1.02% ΔF/F, peak signals during baseline: 1.13 ± 0.24% ΔF/F*, t*(5) = 0.035; *p* = 0.97; unpaired *t*-test*, N* = 6 slices. **E.** Schematic of AAV-CBA-DIO-GCaMP6s injection and fiber implantation followed by discriminatory auditory fear conditioning paradigm with paired (CS⁺) and unpaired (CS^-^) auditory sweeps. **F.** Schematic of fiber photometry setup used to record Ca^2+^ signals in the TeA *in vivo*. **G.** Exemplary epifluorescence image showing *Adcyap1r1*-Cre^+^ neurons expressing GCaMP6s and fiber implant location. **H.** Normalized fluorescence changes (ΔF/F) from *Adcyap1r1*-Cre^+^ neurons expressing GCaMP6s aligned to CS⁺ (peak ΔF/F: 31.07 ± 9.03%) and CS^−^ (peak ΔF/F: 27.48 ± 7.08%) presentations during fear conditioning. The sounds and foot shock are indicated by grey and peach-colored vertical bars (peak ΔF/F after shocks: 65.57 ± 9.36%), *N* = 6 mice. **I.** Same as H, but for retrieval session (peak ΔF/F CS^+^: 26.56 ± 8.45%; peak ΔF/F CS^-^: 24.06 ± 5.71%), *N* = 6 mice. **J.** Quantification of correlation between average Ca^2+^ transients during CS^+^ and freezing level during retrieval session in *Adcyap1r1*-Cre^+^ neurons. Dot plot with linear regression and Pearson correlation, *N* = 6 mice. **K.** Schematic of viral injection of AAV-hSyn-DIO-hM4D_i_-mCherry followed by auditory fear conditioning. CNO or saline was administered 30 min prior to memory retrieval testing. **L.** Overlay of viral injection sites for AAV-hSyn-DIO-hM4D_i_-mCherry in Adcyap1r1-Cre mice in TeA. Color scale indicates percentage of mice with detectable virus expression. **M.** Freezing during retrieval following chemogenetic Gi-GPCR activation of TeA *Adcyap1r1*-Cre^+^ neurons. Left: Quantification of freezing during baseline (CNO: 0.00% [0.00-1.54%], saline: 0.00% [0.00-1.54%]; *U* = 118; *p* = 1, Mann-Whitney *U* test), CS⁺ (CNO: 46.07 ± 4.69%; saline: 42.62 ± 3.81%, *t*(29) = 0.568; *p* = 1, unpaired *t*-test) and CS^-^ (CNO: 39.06 ± 5.89%; saline: 32.94 ± 3.92%, *t*(29) = 0.85; *p* = 0.640, unpaired *t*-test). Right: CS⁺-aligned freezing time course for saline-and CNO-treated groups, *N* = 15 (saline) and 16 (CNO) mice. Data are shown as mean ± SEM or median [IQR].

Since increased PACAP levels and PAC1 genetic alterations are associated with increased risk and symptom severity for PTSD in females (King et al., 2017; Mercer et al., 2016; Ressler et al., 2011), we next asked whether TeA *Adcyap1r1*-Cre^+^ neurons are recruited during auditory fear learning and retrieval *in vivo* in female mice. To this end, we expressed AAV-CBA-DIO-GCaMP6s in TeA of Adcyap1r1-Cre mice and recorded bulk Ca^2+^ transients using fiber photometry as a proxy for neuronal activity. The mice were subjected to auditory discriminatory fear conditioning and subsequent memory retrieval with complex sounds (Fig. 2E-G) – a paradigm that was shown previously to be TeA-dependent (Dalmay et al., 2019; Heimbach et al., 2026). During fear conditioning, *Adcyap1r1*-Cre^+^ neurons showed robust calcium transients aligned with both the conditioned sound (CS^+^) and the unpaired sound (CS^-^), with the largest responses observed following foot shock delivery (Fig. 2H). Direct comparison of shock-aligned fluorescence changes and locomotion (speed) showed no strong correlation, suggesting that shock responses were not attributable to movement artifacts (Extended Data Fig. 2B-D). Across conditioning, CS^+^-evoked transients showed a modest decrease but did not significantly differ between early, middle and late trial blocks, indicating that sensory-evoked recruitment of *Adcyap1r1*-Cre^+^ neurons remained largely stable during acquisition, suggesting lack of novelty encoding and sensory habituation (Extended Data Fig. 2E). Further analysis of potentially behaviorally relevant activity patterns revealed that CS^+^-evoked transients predicted performance on the retrieval day as indicated by a positive correlation between average peak transient amplitude and freezing level (Extended Data Fig. 2F). During retrieval, *Adcyap1r1*-Cre^+^ neurons responded to both CS^+^ and CS^-^presentations (Fig. 2I). However, CS^+^-evoked transients were strongest during the first retrieval trials and decreased after, with a significant reduction from trials 1-5 to trials 6-10 (*p* = 0.037; Extended Data Fig. 2G). Of note, across mice, average CS^+^-evoked calcium transients during retrieval were positively correlated with freezing levels (*r* = 0.976; Fig. 2J), indicating that stronger recruitment of *Adcyap1r1-*Cre*^+^*neurons during both conditioning and retrieval was associated with stronger fear memory expression.

To test whether *Adcyap1r1*-Cre^+^ neuronal activity was required for fear memory expression, we chemogenetically inhibited temporal cortex *Adcyap1r1*-Cre^+^ neurons by expressing AAV-hSyn-DIO-hM4D_i_-mCherry in female Adcyap1r1-Cre mice and activated hM4D_i_ with intraperitoneal (i.p.) injection of CNO (4mg/kg) before retrieval (Fig. 2K-L). Our previous work confirmed that i.p. injection of CNO (4mg/kg) does not affect baseline freezing, CS^+^- or CS^-^-evoked freezing, or discrimination behavior, and that estrous stage of female mice does not significantly affect freezing levels during retrieval (Heimbach et al., 2026). Despite the correlation between CS^+^-evoked activity and freezing levels in our fiber photometric experiments (Fig. 2J), chemogenetic inhibition of *Adcyap1r1*-Cre^+^ neurons did not significantly alter freezing levels during baseline, CS^+^, or CS^-^presentations compared with saline-treated controls (Fig. 2M).

These findings suggest that *Adcyap1r1*-Cre^+^ neuronal activity forms internal representations of fear memory expression but that acute suppression of this population is not sufficient to disrupt fear memory-evoked freezing under these experimental conditions.

### Cortical *Adcyap1r1-*Cre*^+^* neurons show projection-specific PACAP responses

Given that chemogenetic suppression of TeA *Adcyap1r1*-Cre^+^ neurons did not alter freezing levels during retrieval, despite their strong recruitment during memory retrieval (Fig. 2I,M), we wondered whether this was due to the putatively high cell type heterogeneity of this population that might obscure functional contributions of specific neuronal subsets. We therefore first examined which types of long-range-projecting neurons were comprised in the *Adcyap1r1*-Cre*^+^*population, and whether these subsets were differentially modulated by PACAP.

To this end, we performed cell-type-specific anterograde tracing by injecting AAV-EF1α-DIO-hChR2(H134R)-mCherry into the TeA of Adcyap1r1-Cre mice (Fig. 3A). Analysis of axonal labeling revealed widespread projections of *Adcyap1r1*-Cre^+^ TeA neurons to cortical and subcortical regions involved in sensory processing, emotional learning and action selection (Fig. 3B-C, Extended Data Fig. 3A-C). Prominent labeling was observed in the lateral amygdala (LA), posterior striatum (STRp) and contralateral TeA, while additional projections targeted the nucleus accumbens (NAc), medial geniculate complex (MG) and several cortical association areas. These findings confirmed that the TeA *Adcyap1r1*-Cre*^+^* neuronal population is comprised of multiple output neurons.

**Fig. 3:**
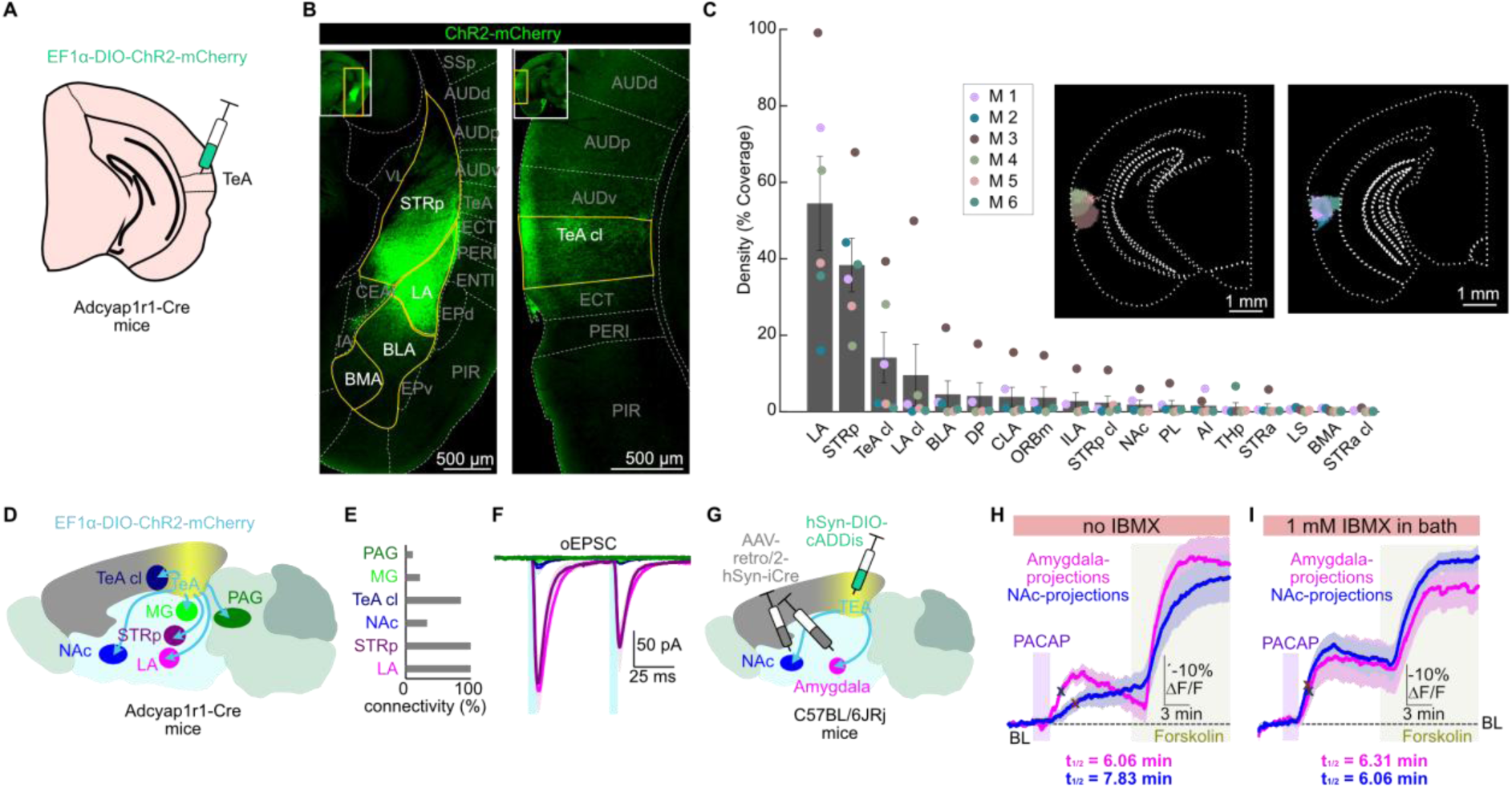
*Adcyap1r1*-Cre^+^ neurons project to multiple cortical and subcortical brain areas and show projection-specific PACAP responses. **A.** Schematic of viral injection of AAV-EF1α-DIO-hChR2(H134R)-mCherry in Adcyap1r1-Cre mice for anterograde tracing of *Adcyap1r1*-Cre^+^ projections from TeA. **B.** Exemplary epifluorescent images of axonal labeling in major target areas. **C.** Overlay of main injection sites of mice used for anterograde tracing and quantification of axonal labeling density per area as percentage of area covered with fluorescent labeling. Each mouse is shown in a different color, *N* = 6 mice. **D.** Schematic of *Adcyap1r1*-Cre^+^ TeA outputs tested in optogenetic-electrophysiological experiments. **E.** Quantification of excitatory connectivity of *Adcyap1r1*-Cre^+^ TeA outputs into indicated target areas in the presence of gabazine and CGP. PAG: 11%, MG: 22%, TeA cl: 85%, NAc: 33%, STRp: 100%, LA: 100%, *N* = 27 cells each. **F.** Average optogenetically evoked excitatory postsynaptic currents (oEPSCs). Same color code as in **E.** Each trace is the mean ± SEM of the connected cells, *N* = 3-27 cells each. **G.** Schematic of viral injections of AAV-hSyn-DIO-cADDis into TeA and AAV-retro/2-hSyn-iCre into amygdala or NAc. **H.** Averaged fluorescence cADDis signals (-ΔF/F) of TeA-amygdala or TeA-NAc projection neurons upon PACAP and forskolin application. Shaded regions indicate ligand application. Peak signals to PACAP: TeA-amygdala projections: -19.81 ± 3.5% ΔF/F; TeA-NAc projections: -14.36 ± 4.53% ΔF/F, *t*(10) = 0.95, *p* = 0.365; unpaired *t*-test. X denotes half-maximum response in TeA-amygdala or TeA-NAc projecting neurons. Amygdala-projecting neurons reached half-maximum significantly earlier than NAc-projecting neurons (TeA-amygdala: 6.06 ± 0.14 min; TeA-NAc: 7.83 ± 0.56 min, *t*(5.62) = −2.92, *p* = 0.029, Welch’s unpaired *t*-test), *N* = 6 slices per condition. **I.** Same as for H, but with IBMX added to the bath. Peak signals to PACAP: TeA-amygdala projections: -37.14 ± 12.93% ΔF/F; TeA-NAc projections: -40.23 ± 7.31% ΔF/F, *t*(10) = 0.21, *p* = 0.84; unpaired *t*-test, *N* = 7-8 slices per condition. Time to half-maximum showed no significant difference between the projections. (TeA-amygdala: 6.62 [6.11-6.85] min; TeA-NAc: 6.0 [5.83 – 6.37] min, *U* = 14, *p* = 0.121, Mann-Whitney *U* test). *N* = 7 slices for TeA-amygdala, *N* = 8 slices for TeA-NAc. **Abbreviations**: AI, agranular insular area; AUD, auditory area; BLA, basolateral amygdala; BMA, basomedial amygdalar nucleus; cl, contralateral; CLA, claustrum; DP, dorsal peduncular area; ECT, ectorhinal area; ENTI, entorhinal area; EP, endopiriform nucleus; ILA, infralimbic area; LA, lateral amygdala; LS, lateral septal nucleus; MG, medial geniculate complex; NAc, nucleus accumbens; ORBm, orbital area, medial part; PAG, periaqueductul gray; PER, perirhinal area; PIR, piriform area; PL, prelimbic area; STRa, anterior striatum; STRp, posterior striatum; SSp, primary somatosensory area; TeA, temporal association area; THp, posterior thalamus, VL, lateral ventricle. Data are shown as mean ± SEM or median [IQR].

We next tested whether anatomically identified projections form functional synaptic connections by combining optogenetic stimulation of *Adcyap1r1*-Cre^+^ TeA axons with whole-cell recordings from neurons in multiple target regions (Fig. 3D). Since a majority of cortical projection neurons are glutamatergic, we blocked GABAergic neurotransmission pharmacologically with gabazine and CGP. Excitatory connectivity was observed in all tested target areas, however, connection probability varied substantially across target areas (Fig. 3E-F). Connectivity was strongest in the LA and STRp, where all recorded neurons responded to photostimulation. High connectivity was also found in contralateral TeA (85% of neurons received glutamatergic input). Lower proportions of connected cells were observed in the NAc (33%), MG (22%) and PAG (11%). Together, these findings demonstrate that *Adcyap1r1*-Cre^+^ TeA neurons establish functional excitatory outputs across a distributed network of sensory, limbic and striatal targets.

Given the putative functional diversity of different *Adcyap1r1*^+^ TeA outputs (Cheng et al., 2025; Dalmay et al., 2019; Day et al., 2007; Li et al., 2026), we wondered whether PACAP effects differed between distinct projection neurons and selected two of the most functionally diverse target areas, the amygdala and NAc. To selectively monitor amygdala-and NAc-projecting neurons, we injected AAV-retro/2-hSyn-iCre into either the amygdala or NAc and expressed the Cre-dependent cAMP sensor in TeA neurons using AAV-hSyn-DIO-cADDis (Fig. 3G). Bath application of 300 nM PACAP evoked significant fluorescence changes, suggesting that cAMP levels increase in both TeA-amygdala and TeA-NAc projection neurons (Fig. 3H, Extended Data Fig. 3D). Peak PACAP-evoked cAMP transients did not differ significantly between TeA-amygdala and TeA-NAc projection neurons (*p* = 0.365). However, TeA-amygdala neurons reached half-maximum significantly earlier than TeA-NAc neurons (*p* = 0.029). We asked whether this apparent difference could reflect different PACAP signaling kinetics or differential shaping of local cAMP dynamics by the enzymatic degradation of cAMP through intracellular phosphodiesterase (PDE) activity. To test this, we repeated the recordings in the presence of the PDE inhibitor IBMX (Fig. 3I). IBMX increased PACAP-evoked fluorescence in both projection neuron populations. Whereas peak amplitudes remained comparable (*p* = 0.84), IBMX eliminated the latency differences between TeA-amygdala and TeA-NAc neurons (*p* = 0.121). Together, our findings indicate that PACAP signaling across distinct *Adcyap1r1*-Cre^+^ projection neurons is shaped by cell type-specific molecular context.

Since PACAP increased intracellular cAMP levels in TeA neurons, and cAMP is a common modulator of synaptic strength (Citri & Malenka, 2008), we next asked whether PACAP acutely modulates synaptic transmission at local or long-range *Adcyap1r1*-Cre^+^ TeA synapses. To this end, we expressed AAV-EF1α-DIO-hChR2(H134R)-mCherry in *Adcyap1r1*-Cre^+^ TeA neurons to optogenetically activate this cell population while recording excitatory postsynaptic currents (EPSCs) from mCherry-negative postsynaptic neurons in the TeA and LA (Extended Data Fig. 3E-F). Neither at local TeA synapses (Extended Data Fig. 3E), nor at TeA-to-LA projections (Extended Data Fig. 3F), did PACAP application significantly alter optogenetically-evoked postsynaptic currents as indicated by unchanged EPSC amplitude and paired-pulse ratios, suggesting that PACAP does not alter presynaptic vesicle release or short-term plasticity. In contrast, artificially increasing cAMP levels with forskolin significantly increased EPSC amplitude (Extended Data Fig. 3G). These findings indicate that, although *Adcyap1r1*-Cre^+^ synapses have the machinery for cAMP-mediated modulation, PACAP does not efficiently engage this signaling mechanism. Consistent with this, PACAP did not significantly alter spontaneous EPSC frequency or amplitude in TeA neurons (Extended Data Fig. 3H).

Together, these experiments show that TeA *Adcyap1r1*-Cre^+^ neurons innervate multiple cortical and subcortical targets and respond to PACAP with projection-specific dynamics of cAMP signaling.

### Cortical Sst^+^ neurons show cell type-specific PACAP signaling and respond to auditory and aversive cues during fear learning and retrieval

Since our expression analysis revealed that cortical *Adcyap1r1* expression is particularly high in GABAergic neurons (Fig. 1), we wondered whether specific inhibitory neuron subclasses could be the main targets of PACAP signaling, thereby increasing feedforward inhibition and limiting the behavioral effects of *Adcyap1r1*-Cre^+^ projection neurons. Hence, we expressed AAV-hSyn-DIO-cADDis in the TeA of Sst-, Pvalb- and Vip-Cre mouse lines and performed epifluorescence cAMP imaging in acute brain slices (Fig. 4A). Bath application of PACAP induced fluorescence transients in Pvalb^+^, Sst^+^, and Vip^+^ neurons, consistent with activation of cAMP signaling (Fig. 4B).

**Fig. 4:**
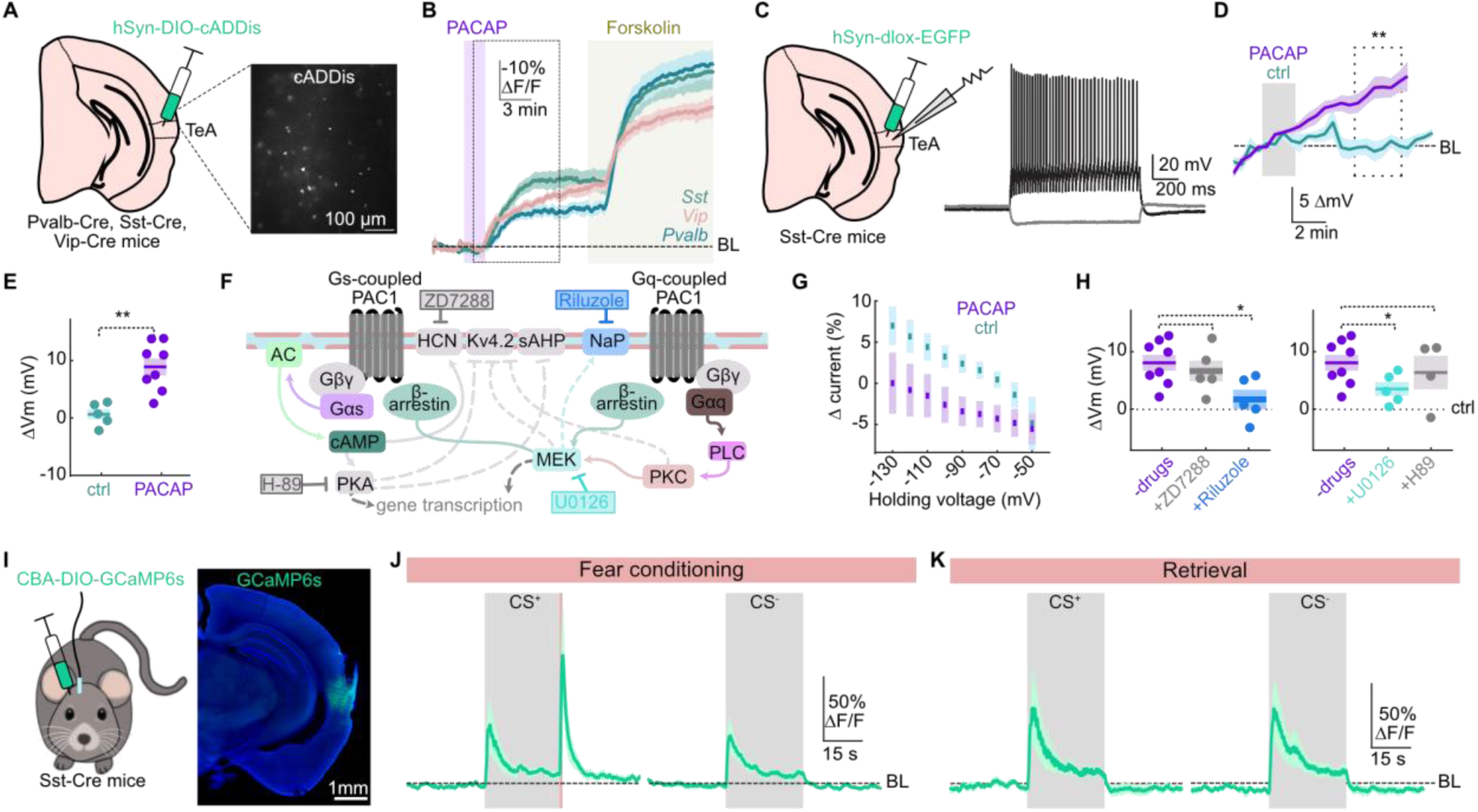
Sst^+^ neurons are recruited by PACAP and respond to auditory and aversive cues during fear learning and retrieval. **A.** Schematic of experimental setup for epifluorescence imaging. AAV-hSyn-DIO-cADDis was expressed in Cre-driver lines in the TeA. Right, exemplary epifluorescent image of cADDis labeling in an acute slice expressed in a Sst-Cre mouse. **B.** Averaged fluorescence cADDis signals (-ΔF/F) of genetically defined neuronal populations following PACAP and forskolin application. Shaded regions indicate drug application. Dashed box indicates area for peak signal analysis. Peak signals to PACAP: Pvalb -26.84% [-22.42--32.9%] ΔF/F; Sst -32.64% [-29.23--61.79%] ΔF/F; Vip -31.73 ± 3.11% ΔF/F. *N* = 10 slices per condition. **C.** Schematic of AAV-hSyn-DIO-EGFP injection in Sst-Cre mice for electrophysiological recordings (left) and exemplary firing pattern of an EGFP^+^ cell in response to hyperpolarizing and depolarizing current injection (right). **D.** Membrane potential changes upon PACAP or vehicle (ctrl) application. Dashed box indicates area for quantification and statistical comparison shown in E. *N* = 5 (ctrl) and 8 (PACAP) cells. BL, baseline. **E.** Quantification for data in **D.**: *t*(11) = -4.08, *p* = 0.002, unpaired *t*-test. **F.** Schematic of signaling cascades downstream of PACAP receptor activation with action sites of the used drugs indicated. Experimentally demonstrated pathways are indicated by colored, solid lines. **G.** Voltage-dependence of PACAP-evoked currents was measured across holding potentials of -130 --50 mV. *N* = 5 (ctrl) and 8 (PACAP) cells. **H.** Quantification of membrane potential changes upon PACAP application in the presence of indicated inhibitors to block HCN channels (ZD7288), persistent sodium channels (riluzole), MEK kinases (U0126) and PKA (H-89) compared to PACAP application in the absence of inhibitors. Unpaired *t*-test for ZD7288: *t*(11) = 0.093, *p* = 0.93; for riluzole: *t*(11) = 3.09, *p* = 0.02; for U0126: *t*(11) = 2.96, *p* = 0.03; for H-89: *t*(10) = 0.81, *p* = 0.44. *N* = 8 (ctrl), 5 (ZD7288), 5 (riluzole), 5 (U0126) and 4 (H-89) cells. **I.** Schematic of the viral injection of AAV-CBA-DIO-GCaMP6s and fiber implantation into the TeA of Sst-Cre mice with exemplary epifluorescence image showing viral expression and fiber implant location. **J.** Normalized fluorescence changes (ΔF/F) from Sst^+^ neurons expressing GCaMP6s aligned to CS⁺ (peak ΔF/F: 51.49 ± 23.85%) and CS^−^ (peak ΔF/F: 41.26 ± 14.81%) presentations during fear conditioning. Peak ΔF/F after shocks: 102.37 ± 40.05%, *N* = 5 mice. **K.** Same as in J., but during fear retrieval (Peak ΔF/F CS^+^: 65.37 ± 30.19%; peak ΔF/F CS^-^: 67.95 ± 26.68%), *N* = 5 mice. **Abbreviations**: AC = adenylyl cyclase; CAV = voltage-gated calcium channels; HCN = hyperpolarization-activated cyclic nucleotide-gated channel; MEK = mitogen activated protein kinase; PKA = protein kinase A; PKC = protein kinase C; PLC = phospholipase C Data are shown as mean ± SEM or median [IQR].

Given that artificial cAMP elevation but not PACAP (Extended Data Fig. 3E-G) enhanced neurotransmission in glutamatergic cells, we wondered whether PACAP induces synaptic modulation in GABAergic neurons. To this end, we expressed AAV-EF1α-DIO-hChR2(H134R)-mCherry in Sst^+^ or Pvalb^+^ TeA neurons to optogenetically activate these GABAergic populations while recording GABAergic postsynaptic currents (PSCs) from nearby mCherry-negative principal cells (Extended Data Fig. 4A-B). Neither Pvalb^+^ nor Sst^+^ synapses onto principal cells were significantly altered by PACAP application (Extended Data Fig. 4A-B), suggesting a primary somatic or dendritic action.

Since accumulating literature suggests that a common intracellular effect of PACAP is depolarization and enhanced excitability (Johnson et al., 2019), we set out to compare these parameters in GABAergic and glutamatergic neurons. To this end, we first expressed AAV-hSyn-DIO-EGFP in the TeA of Sst-Cre mice and performed whole-cell current-clamp recordings of EGFP^+^ cells in acute slices (Fig. 4C). Sst^+^ cell identity was additionally verified by their typical adapting firing pattern with sag potential (Fig. 4C). PACAP application produced a significant depolarization and a trend towards increased excitability (measured by recording the rheobase) compared to vehicle control application (Fig. 4D-E, Extended Data Fig. 4C). In contrast, both depolarization and rheobase changes were non-significant in glutamatergic neurons (Extended Data Fig. 4D). Previous work on other brain areas showed that PACAP-induced depolarization is dependent on pathways involving hyperpolarization-activated cyclic nucleotide-gated channels (HCN) (Sun et al., 2003), persistent NA^+^ currents (NaP) (Johnson et al., 2020), voltage-dependent potassium channels (Kv4.2) (Gupte et al., 2016; Han & Lucero, 2006) or slow afterhyperpolarization (sAHP) potassium conductance (Taylor et al., 2014) (Fig. 4F). To broadly distinguish between potassium and sodium-based conductances, we tested the voltage-relationship of the involved channels using a voltage ramp protocol. PACAP induced a negative shift across the entire voltage range tested (-130 to -50 mV) (Fig. 4G), suggesting involvement of sodium-conducting channels. We therefore pharmacologically blocked HCN and NaP channels with riluzole and ZD7288 respectively. Riluzole, but not ZD7288 significantly reduced PACAP-induced depolarization (Fig. 4H). Further, consistent with previous findings in dentate gyrus neurons (Johnson et al., 2020), pharmacological inhibition of mitogen-activated protein kinase (MEK) with U0126 significantly attenuated depolarization (Fig. 4H), suggesting that MEK-dependent activation of NaP channels contributes to PACAP-induced depolarization in cortical Sst^+^ cells (Fig. 4F). These findings suggest that whereas PACAP signaling affects a broad range of neurons through intracellular cAMP signaling, cAMP-independent pathways favor cortical GABAergic inhibition over direct excitation on the timescales of minutes.

Since Sst^+^ neurons in the cortex provide strong inhibition to apical dendrites of principal cells, thereby reducing the excitatory and plasticity-inducing effect of glutamatergic inputs (Schulz et al., 2018), we next asked whether Sst^+^ neurons in TeA show sensory-and fear memory-related activity in line with inhibitory effects on the mainly glutamatergic *Adcyap1r1*-Cre^+^ neuronal population (Fig. 2H-J). To this end, we expressed AAV-CBA-DIO-GCaMP6s in the TeA of Sst-Cre mice (Fig. 4I) and performed fiber photometry recordings during the same fear conditioning and retrieval paradigm as for Adcyap1r1-Cre mice (Fig. 2E-F). Sst^+^ neurons exhibited calcium transients during CS^+^ and CS^-^presentations during conditioning, as well as large responses following foot shock delivery (Fig. 4J). Similar to *Adcyap1r1*-Cre^+^ neurons, CS^+^ evoked calcium amplitudes in Sst^+^ neurons did not significantly differ among early, middle, and late conditioning trials (Extended Data Fig. 4E). Peak shock-evoked Ca^2+^ transients were negatively correlated with peak locomotor speed during shocks, indicating that shock responses did not represent movement artifacts (Extended Data Fig. 4F-H). During retrieval, Sst^+^ neurons responded robustly to both CS^+^ and CS^-^presentations (Fig. 4K). However, in contrast to *Adcyap1r1*-Cre^+^ neurons, early, middle and late retrieval trials were not significantly different (Extended Data Fig. 4I). Moreover, unlike *Adcyap1r1*-Cre^+^ neurons (Fig. 2J), CS^+^-evoked calcium transients in Sst^+^ neurons during retrieval or conditioning showed no positive correlation with freezing levels during retrieval (retrieval: *r* = -0.056; conditioning: *r* = 0.110; Extended Data Fig. 4J-K). To test whether the lack in correlation could be due to high variability in CS^+^ responses between mice, we normalized retrieval CS^+^ responses to conditioning CS^+^ responses. Indeed, when comparing normalized CS^+^-evoked calcium transients during retrieval to freezing levels during retrieval, we observed a more pronounced negative correlation (*r* = -0.492; Extended Data Fig. 4L). This was in contrast to the positive correlation observed when performing the same operations for *Adcyap1r1*-Cre^+^ neuron recordings (*r* = 0.682; Extended Data Fig. 4M). Thus, our data reveal that, although basic sound and shock responses are present in both Sst^+^ and *Adcyap1r1*-Cre^+^ neurons, more complex computations, including sound habituation and correlations between sound-evoked activity and freezing, exhibit opposing features.

Together, our findings identify a PACAP signaling pathway that favors depolarization in inhibitory but not excitatory TeA neurons, thereby facilitating differential modulation of functionally distinct cell types despite broad receptor expression. Moreover, the distinct sensory- and fear-related activity patterns of inhibitory Sst^+^ neurons *in vivo* support a circuit model in which these neurons modulate dendritic inhibition of principal cells in a computation-specific manner, thereby shaping TeA output during fear-memory processing.

### PACAP engages Gq-dependent Ca^2+^ signaling and cAMP signaling in TeA astrocytes

Our expression analysis revealed high *Adcyap1r1* expression in TeA astrocytes (Fig. 1), and our previous work showed that, in contrast to neuronal populations, astrocytes respond to PACAP with robust Ca^2+^ elevations and that Ca^2+^ dynamics in cortical astrocytes are essential for fear memory retrieval (Heimbach et al., 2026). We therefore hypothesized that PACAP recruits astrocyte-specific functionally relevant intracellular pathways. To investigate the molecular pathways underlying PACAP-evoked astrocytic Ca^2+^ signaling in detail, we expressed AAV-hGFAP-GCaMP6f in TeA astrocytes and pharmacologically dissected candidate pathways downstream of GPCR activation in acute slices (Fig. 5A-D). PACAP induced robust Ca^2+^ transients under control conditions, whereas blockade of Gq signaling with YM-254890 significantly reduced the response (*p* = 0.011). Similarly, inhibition of IP_3_ receptors with 2-APB (*p* = 0.018) and inhibition of phosphatidylinositol-specific PLC with U73122 (*p* = 0.020) significantly reduced PACAP-evoked Ca^2+^ transients (Fig. 5B,C). In contrast, inhibition of Gβγ signaling with gallein, L-type voltage-activated Ca^2+^ channels with nimodipine, phosphatidylcholine-specific PLC with D609, or PKA with H-89 did not significantly alter peak PACAP-evoked Ca^2+^ transients compared with control conditions (Fig. 5C; Extended Data Fig. 5A). Moreover, direct activation of adenylyl cyclase with forskolin did not evoke a significant astrocytic Ca^2+^ transient in comparison with control (*p* = 0.511; Extended Data Fig. 5B). Together, these data indicate that PACAP-evoked astrocytic Ca^2+^ elevations are primarily mediated through a Gq/PLC/IP_3_-dependent pathway rather than through cAMP/PKA signaling or voltage-gated Ca^2+^ entry (Fig. 5D). Moreover, application of the selective PAC1 agonist maxadilan evoked similar Ca^2+^ elevations in comparison to PACAP (*p* = 0.529, Fig. 5E), indicating that, in astrocytes, PACAP primarily binds PAC1 rather than the lower affinity receptors VPAC1/2.

**Fig. 5:**
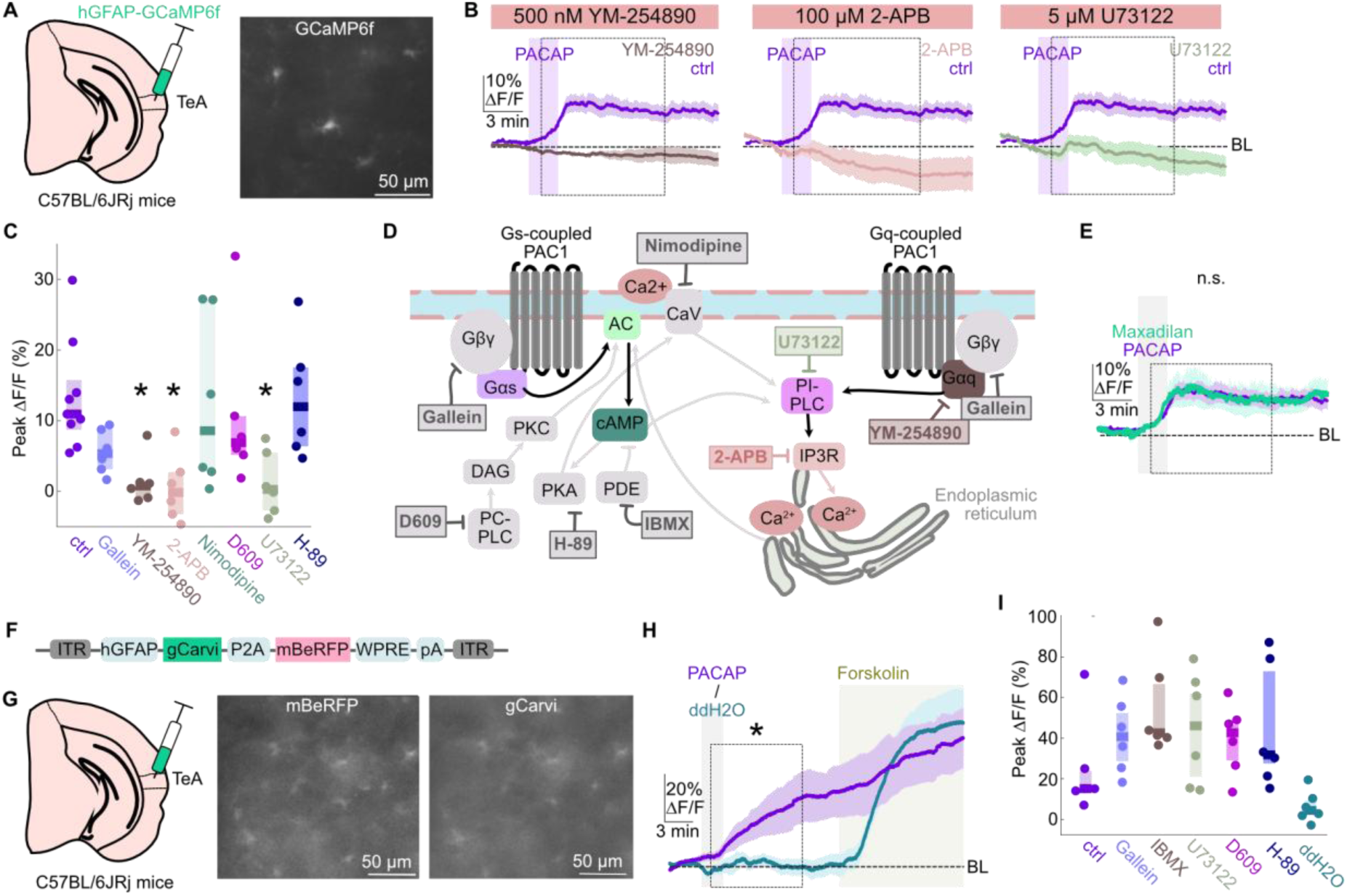
PACAP engages Gq-dependent Ca^2+^ signaling and cAMP signaling in TeA astrocytes. **A.** Schematic of viral injection of AAV-hGFAP-GCaMP6f into TeA and exemplary epifluorescence image of GCaMP6f labeling in an acute TeA slice. **B.** Averaged fluorescence Ca^2+^ signals (ΔF/F) from TeA astrocytes in response to PACAP application with YM-254890, 2-APB and U73122 supplemented to the bath compared to controls. Purple boxes indicate PACAP application. *N* = 6 -8 slices per condition. **C.** Averaged peak fluorescence Ca^2+^ signals (ΔF/F) from TeA astrocytes in response to PACAP application in the absence (ctrl) and presence of indicated blockers. Peak signals to PACAP: ctrl: 13.37 ± 2.58% ΔF/F; 10.93 % [8.71-15.75%] ΔF/F; gallein: 5.6 ± 1.26% ΔF/F, *t*(13) = 2.32, *p* = 0.149; unpaired *t*-test; YM-254890: 0.75% [-0.91-1.27%] ΔF/F, *U* = 2, *p* = 0.011; Mann-Whitney *U* test; 2-APB: 0.48 ± 1.94% ΔF/F, *t*(13) = 3.632, *p* = 0.018; unpaired *t*-test; nimodipine: 12.43 ± 5.02% ΔF/F, *t*(13) = 0.184, *p* = 1; unpaired *t*-test; D609: 6.88% [5.15-10.65%] ΔF/F, *U* = 16, *p* = 0.671; Mann-Whitney *U* test; U73122: 1.15 ± 1.83% ΔF/F, *t*(13) = 3.477, *p* = 0.02; unpaired *t*-test; H-89: 13.23 ± 3.43% ΔF/F, *t*(13) = 0.035, *p* = 1; unpaired *t*-test; *N* = 6-9 slices per condition. **D.** Schematic of signaling cascades downstream of GPCR activation in astrocytes with action sites of drugs used in C and H indicated. Experimentally suggested pathways represented by colored arrows. **E.** Averaged fluorescence Ca^2+^ signals (ΔF/F) from TeA astrocytes in response to PACAP and maxadilan (150 nM) application. Maxadilan peak signals: 6.37% [4.24-8.4%] ΔF/F, statistical comparison to PACAP: *U* = 21, *p* = 0.529, Mann-Whitney *U* test. *N* = 9 (PACAP) and 6 (maxadilan) slices. **F.** Schematic of viral construct for astrocytic cAMP imaging. **G.** Left: Schematic of viral injection for TeA astrocytic cAMP imaging experiment. Right: Exemplary epifluorescence image of mBeRFP and gCarvi labeling in an acute TeA slice. **H.** Averaged fluorescence cAMP signals (ΔF/F) from TeA astrocytes in response to PACAP and forskolin application compared to vehicle (ddH2O). Peak signals: PACAP: 21.19% [17.17-25.32%] ΔF/F; control: 6.87% [1.74-14.63%] ΔF/F; *U* = 3; *p* = 0.015, Mann-Whitney *U* test, *N* = 6 slices per condition. **I.** Averaged peak fluorescence cAMP signals (ΔF/F) from TeA astrocytes in response to PACAP in the absence (ctrl) and presence of indicated blockers. Peak signals PACAP control: 21.19% [17.17-25.32%]; gallein: 40.71% [25.42-55.64%] ΔF/F, *U* = 7, *p* = 0.279; IBMX: 42.64% [40.32-69.79%] ΔF/F, *U* = 5, *p* = 0.206; U73122: 46.07 % [15.43-67.55%] ΔF/F, *U* = 9, *p* = 0.359; D609: 42.67% [26.62-48.98%] ΔF/F, *U* = 10, *p* = 0.359; H-89: 31.74% [21.32-79.17%] ΔF/F, *U* = 6, *p* = 0.26; ddH2O control: 6.87% [1.74-14.63%]; Mann-Whitney *U* tests were performed for all conditions; *N* = 6 slices per condition. **Abbreviations**: AC = adenylyl cyclase; CAV = voltage-gated calcium channels; DAG = diacylglycerol; IP3R = Inositol 1,4,5-trisphosphate receptor; PDE = phosphodiesterase; PKA = protein kinase A; PKC = protein kinase C; PC-PLC = phosphatidylcholine-specific phospholipase C; PI-PLC = phosphoinositide-specific phospholipase C. Data are shown as mean ± SEM or median [IQR].

To further investigate the involvement of astrocytic Gq signaling, we expressed iβARK, a genetically encoded tool developed to attenuate astrocytic Gq-GPCR signaling, in TeA astrocytes (Nagai et al., 2021) (Extended Data Fig. 5C-D). However, iβARK expression did not significantly reduce astrocytic Ca^2+^ transients evoked by either PACAP or norepinephrine compared with the inactive iβARK(D110A) mutant control (Extended Data Fig. 5E-F). Consistent with previous reports that iβARK-mediated inhibition of Gq-GPCR signaling can be incomplete under certain conditions (Nagai et al., 2021), we interpret our findings as reflecting context-dependent efficacy of iβARK (for example due to differences in receptor localization or abundance) rather than as evidence for Gq-independent Ca^2+^ signaling in TeA astrocytes.

We next set out to test whether PACAP also regulates cAMP signaling in astrocytes. Since no green fluorescent cAMP sensor specifically suited for astrocytes was available, we developed a new genetically encoded cAMP sensor construct based on gCarvi (Kawata et al., 2022), under the control of the hGFAP promoter, and comprising a P2A-linked long-stokes shifted red fluorescent protein, mBeRFP for identification of gCarvi expressing-cells independently of astrocytic cAMP fluctuations (Fig. 5F). We injected the newly generated AAV-hGFAP-gCarvi-P2A-mBeRFP into the TeA. gCarvi and mBeRFP fluorescence were detected in cells with typical astrocytic morphology (Fig. 5G). PACAP induced significant astrocytic gCarvi transients compared with vehicle application (*p* = 0.015; Fig. 5H), demonstrating that PACAP can engage both Ca^2+^ and cAMP signaling in cortical astrocytes, which is in line with previous studies showing PACAP-induced cAMP production and PLC/IP_3_ signaling in cultured astrocytes (Dejda et al., 2006; Masmoudi et al., 2003; Sauer et al., 2026).

Because cAMP signaling can be modulated by Ca^2+^ through Ca^2+^/calmodulin-sensitive adenylate cyclase isoforms (Sauer et al., 2026), U73122 was used to test whether PLC/ Ca^2+^-dependent signaling contributed to the cAMP elevations. Of note, inhibition of PLC with U73122, which significantly reduced PACAP-evoked Ca^2+^ transients (Fig. 5C), did not detectably affect peak cAMP transients (*p* = 0.359, Fig. 5I, Extended Data Fig. 5G). We further tested potential Gβγ-dependent modulation of adenylyl cyclase activity with gallein (Hanoune & Defer, 2001), PKA-dependent feedback with H-89 (Desrues et al., 2008), and PDE-mediated cAMP degradation with IBMX (Christiansen et al., 2011). D609 was additionally used to probe phosphatidylcholine-specific PLC sensitive lipid signaling. None of these manipulations significantly altered the peak PACAP-evoked cAMP transients compared with control conditions (Fig. 5I; Extended Data Fig. 5G). These findings are therefore consistent with the recruitment of distinct intracellular signaling pathways for cAMP and Ca^2+^ transients. One possible mechanism is Gs-dependent activation of adenylyl cyclase in parallel with Gq/PLC/IP_3_-dependent Ca^2+^mobilization. However, the G protein coupling underlying astrocytic cAMP production remains to be determined.

Of note, selective activation of PAC1 with maxadilan evoked large cAMP transients, exceeding those induced by PACAP (*p* = 0.026; Extended Data Fig. 5H), supporting PAC1 as the main receptor mediating PACAP-evoked cAMP signaling in cortical astrocytes.

Together, our findings reveal a cell-type-specific organization of PACAP signaling in the TeA. Whereas PACAP recruits cAMP but not Ca^2+^ signaling in neuronal populations, astrocytes engage both cAMP and Gq-dependent Ca^2+^ signaling. Thus, this further supports the idea that PACAP differentially fine-tunes functionally distinct cell types despite broad receptor expression.

## Discussion

PACAP is an important modulator of neuronal circuits across multiple brain regions and functions (Kirry et al., 2018; Meloni et al., 2016; Schmidt et al., 2015). Its receptors are among the most widely expressed neuropeptide receptors in the cortex (Smith et al., 2019; Zhang et al., 2021). However, the broad distribution of PACAP receptors across cortical cell types with opposing functional roles has remained puzzling (Zhang et al., 2021). Here, we identify a cell-type specific organization of PACAP signaling in the cortex that extends substantially beyond the mere distribution of its receptors. PAC1 receptor expression is widespread across excitatory and inhibitory neurons and non-neuronal cells, particularly astrocytes. Yet, our data reveal that PACAP does not elicit a uniform intracellular response across these populations, but instead engages distinct, cell-type-specific signaling programs. Thus, we provide evidence, that PACAP recruits cAMP-dependent, but not Ca^2+^-dependent pathways in neurons. In astrocytes, by contrast, PACAP recruits both cAMP and Ca^2+^-dependent pathways. Additionally, whereas cAMP is recruited across multiple neuronal cell types, we show that additional cAMP-independent pathways are recruited in a subtype-specific pattern. In particular, PACAP increases the resting membrane potential of Sst^+^ neurons through a MEK-NaP-dependent mechanism within minutes of receptor activation, without producing detectable acute physiological changes in glutamatergic neurons. Together, these findings highlight that in order to predict and understand PACAP effects, detailed cell type-specific analysis of intracellular and physiological responses is required.

A potential basis for these cell-type specific signaling pathways lies in differential PAC1 receptor coupling to G proteins. PAC1 can activate both Gs-adenylyl cyclase-cAMP as well as Gq-PLC-Ca^2+^ signaling, and alternative splicing generates receptor isoforms with distinct efficiencies for activating these downstream cascades (Hammack & May, 2015; Pisegna & Wank, 1996; Spengler et al., 1993). The rodent cortex expresses the canonical PAC1 isoform as well as hip- and hop-containing variants, although the latter occur at lower abundance (Spengler et al., 1993). Whether these variants are differentially distributed between cortical neurons and astrocytes remains unknown. Alternatively, differences in G-protein availability, binding of accessory proteins, receptor localization or downstream signaling machinery may determine which pathway predominates in individual cell types. Our findings that *Adcyap1r1* expression alone does not predict the functional consequence of PACAP signaling underscores the importance of further characterizing cell-type-specific receptor isoform distribution and downstream coupling also for other neuropeptides, cell types and brain areas. Importantly, differential expression of isoforms with differential downstream coupling provides promising opportunities for the development of cell-type-selective pharmacological interventions, enabling more precise modulation of PACAP signaling while minimizing side-effects.

Beyond differences in second-messenger recruitment, our findings show that PACAP signaling can induce cell-type specific changes in intrinsic neuronal excitability. In Sst^+^ neurons, PACAP induced a progressive depolarization that was dependent on the recruitment of MEK-dependent persistent Na^+^ conductance. This resembles PACAP signaling described in dentate gyrus granule cells (Johnson et al., 2020). One potential mechanism linking PAC1 activation to this relatively slow change is β-arrestin-dependent receptor internalization and endosomal MEK/ERK signaling. PAC1 can recruit β-arrestins, which in turn have been implicated in PAC1 internalization and ERK1/2 activation (Shintani et al., 2018). Moreover, inhibition of PAC1 internalization or MEK/ERK signaling attenuates PACAP-induced increases in neuronal excitability in other neuronal populations (May et al., 2021). Thus, β-arrestin-dependent endosomal signaling represents a candidate mechanism for the PACAP-induced depolarization in Sst^+^ neurons.

The astrocytic PACAP-evoked signaling cascades represent a particularly novel insight into the diversity of PACAP effects. Whereas our recent work firstly identified astrocytes as a target of PACAP signaling in TeA (Heimbach et al., 2026), another recent study revealed astrocytic L-type Ca^2+^ channels as main mediators of PACAP-induced Ca^2+^ transients in prefrontal cortex (Cui et al., 2025). Surprisingly, our new findings show that PACAP does not only have cell type-specific but also cortical area-specific signaling effects. Thus, in TeA astrocytes, PACAP recruits intracellular Ca^2+^ in an endoplasmatic reticulum-dependent and L-type Ca^2+^ channel-independent manner. This is fundamentally different from the signaling pathway discovered in prefrontal cortex astrocytes. In addition, TeA astrocytes increase intracellular cAMP levels following PACAP application, showing that PACAP can recruit multiple intracellular pathways within the same cell type. Whether these different PACAP-evoked astrocytic pathways result in divergent or shared downstream network effects, such as gliotransmission, potassium buffering and synaptic modulation, and whether the two signaling pathways result from differential isoform expression, remain important questions for future research.

Our *in vivo* findings provide functional insights into cell type-specific behavioral representations of target cells, thereby providing a framework for how PACAP could modulate fear processing in the TeA. Thus, we provide evidence that *Adcyap1r1*-Cre^+^ TeA neurons are strongly recruited by auditory cues and foot shock during fear conditioning and retrieval, and CS^+^-evoked activity correlates with freezing. In contrast, Sst^+^ neurons respond similarly to sounds and shocks, but their activity during CS^+^ is negatively correlated with fear-memory expression. Of note, whereas the *Adcyap1r1*-Cre^+^ population also includes Sst^+^ cells, the bulk photometry signals are likely dominated by excitatory neurons due to their high abundance and high surface area in the cortex. The opposing correlations between calcium dynamic in glutamatergic and Sst^+^ neurons and fear expression are consistent with a circuit model in which Sst^+^ neurons preferentially target the dendritic compartments of glutamatergic neurons, thereby regulating dendritic integration and NMDA receptor-dependent excitation (Lovett-Barron et al., 2014; Schulz et al., 2018). In the hippocampus, recruitment of dendrite-targeting interneurons during aversive stimulation limits dendritic excitation and is required for fear learning (Lovett-Barron et al., 2014). PACAP-dependent elevation of Sst^+^ membrane potential could therefore increase the probability of Sst^+^ recruitment when excitatory drive is present, thereby increasing dendritic inhibition, even in the absence of a direct PACAP-induced presynaptic change in transmitter release at Sst^+^ synapses. Based on our fiber photometric recordings showing opposing correlations of neuronal activity and freezing, we hypothesize that increased Sst^+^ neuron activity in TeA attenuates inputs to glutamatergic neurons carrying learned and behaviorally relevant information (conditioned sounds), while having less influence on neutral sensory inputs. This could explain why neuronal *Adcyap1r1*-Cre^+^ and Sst^+^ populations both respond strongly to auditory and aversive stimuli while showing partially opposing relationships with fear-memory expression.

Of note, whereas our photometry results suggest that *Adcyap1r1*-Cre^+^ and Sst^+^ neurons encode sensory inputs and behavioral outcomes, behaviorally neither cell type had an effect on fear memory expression after chemogenetic inhibition (Heimbach et al., 2026). In the case of *Adcyap1r1*-Cre^+^ neurons, simultaneous chemogenetic inhibition of excitatory and inhibitory neuronal populations may produce opposing circuit effects, potentially obscuring the functional contributions of individual *Adcyap1r1*-Cre^+^ neuronal subtypes to fear-memory retrieval. Whereas inhibitory neurons represent only ∼20% of the neuronal population (Tremblay et al., 2016), they can have substantial functional effects due to their extensive axonal arborization and strong inhibition of surrounding principal cells (Lovett-Barron et al., 2014; Schulz et al., 2018; Urban-Ciecko & Barth, 2016). Moreover, inhibition of *Adcyap1r1*-Cre^+^ neurons may simultaneously affect different TeA projection populations with different, or potentially opposing, behavioral functions. Indeed, different TeA output pathways have been implicated in distinct aspects of defensive behavior. Thus, projections to the lateral amygdala and tail of the striatum contribute to auditory threat memory (Cheng et al., 2025; Dalmay et al., 2019), whereas TeA layer 5 neurons projecting to the dorsal periaqueductal gray drive sensory-induced escape behavior (Li et al., 2026). Future cell type- and projection-specific manipulations will be required to distinguish the functional contributions of PACAP-PAC1 signaling in these cell types for fear memory expression.

Alternatively, *Adcyap1r1*-Cre^+^ neurons in TeA may reflect internal representations of sensory cues and their learned emotional value rather than directly contributing to the overall motor expression of freezing. Previous studies support the view that cortical neuronal activity and its functional effects are highly context-specific, without having broad effects on behavioral output. Thus, Russell et al. demonstrated functional effects under conditions of high task engagement (Russell et al., 2024). In line with this, the strong correlation between *Adcyap1r1*-Cre^+^ neuron activity and freezing levels may reflect an internal cortical representation associated with fear-memory expression that becomes functionally relevant under specific task conditions, while other circuits determine whether and how this representation is translated into overall freezing behavior under our experimental conditions. This fits well with our own recent work showing that manipulation of TeA astrocytes but not TeA neurons alters fear memory expression (Heimbach et al., 2026). Future research will reveal whether TeA *Adcyap1r1*-Cre^+^ neurons and their activity patterns contribute to context-dependent behavioral outputs or are more relevant for other aspects of fear-related behavior, such as remote fear memory retrieval or escape behavior.

PACAP-PAC1 signaling is particularly well established in the context of stress and fear, with studies implicating this peptidergic signaling channel across several brain regions. PACAP signaling in the amygdala, hippocampus and prefrontal cortex can modulate stress responses, synaptic plasticity and aversive learning (Hirabayashi et al., 2018; Kirry et al., 2018; Meloni et al., 2016; Rajbhandari et al., 2021; Schmidt et al., 2015). Moreover, human studies have associated circulating PACAP levels and *ADCYAP1R1* variation with PTSD symptoms, with particularly prominent associations reported in females (King et al., 2017; Mercer et al., 2016; Ressler et al., 2011). Beyond stress-related disorders, altered PACAP signaling has been implicated in neurodegenerative disease, including Alzheimer’s disease (Basri et al., 2026) and Parkinson’s disease (Hassan et al., 2026), while PACAP and its receptors have received considerable attention as therapeutic targets in migraine (Amin et al., 2014). Together, these observations emphasize the need to understand how PACAP acts within intact and pathological brain circuits. Our findings show that the consequences of PACAP receptor activation cannot be inferred from receptor expression alone, but depend strongly on the identity and intracellular signaling architecture of the responding cell.

Thus, the question arises, as to whether endogenously released PACAP can reach different cell types with different spatio-temporal concentration profiles. Whereas *Adcyap1r1* expression is widespread, our data show that *Adcyap1* expression is comparatively restricted and enriched in subsets of cortical glutamatergic neurons. Localized PACAP release could therefore recruit different combinations of receptor-expressing neurons and astrocytes and, through their distinct intracellular and circuit functions, differentially alter cortical processing. In such a scenario, the net effect of endogenous PACAP release would depend strongly on the cellular populations reached by the released peptide. Consequently, a major limitation is that direct monitoring of endogenous PACAP release remains technically challenging, highlighting the importance of further optimizing PACAP sensors with high temporal and spatial resolution (Cola et al., 2024).

Together, our findings reveal a highly cell type- and brain area-specific PACAP signaling architecture across excitatory and inhibitory neuronal populations and astrocytes. We demonstrate that the functional consequences of PACAP cannot be inferred from PACAP receptor expression alone and instead depend strongly on cellular identity and signaling context. Such diversification provides a mechanism through which a single neuropeptide can differentially modulate functionally distinct cell types in cortical circuits involved in emotional learning despite ubiquitous receptor expression.

## Methods

### Animals

The following mouse lines were used in this study: C57BL/6JRj (Janvier; RRID:IMSR_RJ:C57BL-6JRJ), Adcyap1r1-Cre mice (The Jackson Laboratory; RRID: IMSR_JAX:035572), Sst-Cre (B6J.Cg-Ssttm2.1(cre)Zjh/MwarJ; The Jackson Laboratory, RRID:IMSR_JAX:028864), Vip-Cre (B6J.Cg-VIPtm1(cre)Zjh/AreckJ; The Jackson Laboratory, RRID:IMSR_JAX:031628), and Pvalb-Cre (B6.129P2-Pvalbtm1(cre)Arbr/J; The Jackson Laboratory, RRID:IMSR_JAX:017320). All transgenic lines were backcrossed to C57BL/6JRj mice for at least 7 generations. Mice were kept in standard enriched laboratory cages under a 12h light/dark cycle, with *ad libitum* access to food and water. The mice used for *in vitro* experiments were group housed, whereas mice undergoing behavioral experiments were single-housed starting at the second day of handling. Female mice were used for behavioral experiments. Mice of both sexes were used for other experiments.

All procedures were performed in accordance with the institutional guidelines of Medical University of Vienna and approved by the Austrian Ministry of Science.

### Stereotactic surgeries

Mice underwent stereotaxic surgery at 6-12 weeks of age. Anesthesia was induced using 5% isoflurane. Throughout the surgical procedure, mice were kept under stable anesthesia with 1%-2.5% isoflurane and placed on a feedback-controlled heating pad, maintaining a body temperature of 37 °C. Lidocaine (3.5 mg/kg^-1^; Gebro Pharma, cat. no. 100562) and Carprofen (4 mg/kg^-1^; Zoetis, cat. no. 256684) were injected subcutaneously for local anesthesia and general analgesia prior to incision. The skull was leveled using bregma and lambda as reference points (David Kopf Instruments).

A small craniotomy was made at the respective coordinate relative to bregma. For targeting the TeA, injections were performed at -3.6 mm posterior (AP) relative to bregma, 0.35 mm medial to the lateral skull ridge (ML) and -1.23 mm dorsoventral (DV) from the pial surface. For targeting the NAc, injections were made at +1 mm AP, 2.0 mm ML from the midline, and -3.8 mm DV using a 15 ° angle of the injector towards the midline. For targeting the amygdala, injections were made at 0 mm AP, 3.9 mm ML from the midline, and -4.4 mm DV using a 25° posterior angle and 20° lateral rotation of the injector.

For viral injections, AAVs were administered using a glass micropipette at a rate of 40 nL/minute, with injected volumes ranging from 50-200 nL. For chemogenetic manipulation of neurons in behavioral experiments, Adcyap1r1-Cre mice received bilateral injections of 50-100 nL of AAV-9/2-hSyn-DIO-hM4D_i_-mCherry(rev)-WPRE-hGHp(A) (7.7-12*10^11 gc/mL; VVF UZH, cat. no. v84-9 (Krashes et al., 2011) into the TeA. For fiber photometry experiments, 200 nL of AAV-CBA-DIO-GCaMP6s-P2A-mBeRFP-WPRE-pA (2.9*10^12 gc/mL; VVF UZH, cat. no. v828-9) was injected unilaterally into Adcyap1r1-Cre mice or Sst-Cre mice.

For slice imaging experiments targeting astrocytes, 200 nL of AAV-5/2-hGFAP-hHBbl/E-GCaMP6f-bGHp(A) (5.6*10^12 gc/mL; VVF UZH, cat. no. v275-5) or 200 nL of AAV-5/2-hGFAP-gCarvi-P2A-mBeRFP-WPRE-pA (4.5*10^12 gc/mL; VVF UZH, cat. no. vSME25-5) were injected bilaterally into the TeA of C57BL/6JRj mice. For slice imaging experiments targeting neurons, 200 nL of the following AAVs were injected into the TeA of Adcyap1r1-Cre mice, Sst-Cre, Vip-Cre, and Pvalb-Cre mice: AAV-hSyn-DIO-GreenDownward-cADDis (2*10^13 gc/mL; VectorBuilder via Molecular Montana) or AAV-CBA-DIO-GCaMP6s-P2A-mBeRFP-WPRE-pA. For experiments targeting projection neurons, 200nL AAV-hSyn-DIO-GreenDownward-cADDis was injected into the TeA along with 200 nL AAV-retro/2-hSyn-chl-iCre-WPRE-SV40p(A) (7.9*10^12 gc/mL; VVF UZH, cat. no. v223-retro) mixed with AAV-9/2-hSyn-DIO-mCherry(rev)-DIO-WPRE-hGHp(A) (1*10^12 gc/mL; VVF UZH, cat. no. v116-9) to verify the injection site in the NAc or amygdala in C57BL/6JRj mice. For experiments attempting to attenuate Gq-GPCR signaling in astrocytes, 200 nL of AAV-5/2-hGFAP-chI-bBARK(rgs)_2A_mCherry-WPRE-SV40p(A) (986 bp long promoter) (4*10^11 gc/mL, VVF UZH, cat. no. v748-5) or 100 nL of AAV-5/2-hGFAP-chI-bBARK(rgs)(D110A)_2A_mCherry-WPRE-SV40p(A) (986 bp long promoter) (3*10^12 gc/mL, VVF UZH, cat. no. v749-5) were injected into TeA.

For electrophysiology and anterograde tracing experiments, 30-50 nL of AAV-2/2-EF1a-DIO-hChR2(H134R)-mCherry-WPRE-hGHp(A) (3.1*10^12 gc/mL; UNC vector core) or 200 nL of AAV-9/2-hSyn-DIO-EGFP(rev)-WPRE-hGHp(A) (1.6*10^12 gc/mL; VVF UZH, cat. no. v115-9) were injected into the TeA of Adcyap1r1-Cre mice, Pvalb-Cre or Sst-Cre mice.

Before and after the injection, the pipette was kept in place for a wait time of 3 min to minimize reflux. For fiber photometry experiments, the scalp skin was removed and the skull surface was roughened. Mono fiberoptic cannulae (Doric Lenses, cat. no. MFC_200/230–0.37_3mm_MF1.25_A45) were implanted at the injection site of the TeA at -1.23 DV, with the 45° angled fiber tip facing anterior. The craniotomy was sealed with Kwik-cast (World Precision Instruments), and the implant was secured with cyanoacrylate glue reinforced with Paladur (Kulzer, cat. no. L010131) and covered with black nail polish to reduce ambient light scatter.

For the other mice, the skin of the scalp was sutured. The mice were allowed to recover in a heated recovery cage. Post-surgery, mice were monitored for 4 consecutive days, with Carprofen (4 mg/kg) being administered for the first 2 days post-surgery. Following the surgery, a period of 3-4 weeks passed before starting the experiments, allowing for efficient viral expression.

### Plasmids

pAAV-hSyn-DIO-hM4D_i_-mCherry was a gift from Bryan Roth (Addgene plasmid cat. no. 44362; http://n2t.net/addgene:44362; RRID:Addgene_44362).

pAAV-hGFAP-hHBbl/E-GCaMP6f-bGHp(A) was constructed by the VVF (GCaMP6f: Addgene cat. no. 51083; hGFAP promoter fragment: DOI: 10.1002/glia.20622).

pAAV-CBA-DIO-GCaMP6s-P2A-mBeRFP-WPRE-pA was a gift from Bernardo Sabatini (Addgene plasmid cat. no. 175177; http://n2t.net/addgene:175177; RRID:Addgene_175177).

pAAV-hSyn-DIO-GreenDownward-cADDis was purchased from Montana Molecular (Tewson et al., 2016).

pAAV-hSyn-chl-iCre-WPRE-SV40p(A) was constructed by the VVF.

pAAV-EF1a-DIO-hChR2(H134R)-mCherry-WPRE-HGHpA was a gift from Karl Deisseroth (Addgene plasmid cat. no. 20297 ; http://n2t.net/addgene:20297 ; RRID:Addgene_20297).

pAAV-hGFAP-chI-bBARK(rgs)-P2A-mCherry-WPRE-SV40p(A) was a gift from Baljit Khakh (Addgene plasmid cat. no. 117691 ; http://n2t.net/addgene:117691 ; RRID:Addgene_117691). Published in Nagai et al. (2021).

pAAV-hGFAP-chI-bBARK(rgs)(D110A)-P2A-mCherry-WPRE-SV40p(A) was a gift from Baljit Khakh (Addgene plasmid cat. no. 117692 ; http://n2t.net/addgene:117692 ; RRID:Addgene_117692) Published in Nagai et al. (2021).

For the generation of the pAAV-Gfap-gCarvi-P2A-mBeRFP-WPRE-pA plasmid, the sequence GTCGAC in plasmid pAAV-GFAP-hM3D(Gq)-mCherry was changed to ACCGGT to introduce an Age1 restriction site. The plasmid was then cut with restriction enzymes Age1 and EcoRI and the following Kozak-gCarvi-P2A-mBeRFP sequence was inserted via its Age1 and EcoR1 restriction sites: ACCGGTgccaccatggactacaaagaccatgacggtgattataaagatcatgacatcgattacaaggatgacgatgacaagggatccatggtgctgggcaaaccacagaccgacccaacattggagtggtttctgtcacactgtcacattcacaagtacccatctaagagcaccctgatccatcagggcgagaaggctgaaaccctgtattatatagtgaagggttcagtcgccgtgctgattaaggatgaagaaggaaaggaaatgatcctgagctatctgaaccagggtgactttatcggagaactgggcctgttcgaggaaggacaggagagatccgcttgggtgagggctaagacagcctgtgaggtggccgaaatctcatacaagaaattccggcaactcatacaggtgaatcccgacatactcatgcgcctctcagcccagatggcgagaaggcttcaggttacctctgagaaggtggggaacctgtacctcgagaacgtctatatcaaggccgacaagcagaagaacggcatcaaggcgaacttcaagatccgccacaacatcgaggacggcggcgtgcagctcgcctaccactaccagcagaacacccccatcggcgacggccccgtgctgctgcccgacaaccactacctgagcgtgcagtccattctttcgaaagaccccaacgagaagcgcgatcacatggtcctgctggagttcgtgaccgccgccgggatcactctcggcatggacgagctgtacaagggcggtaccggagggagcatggtgagcaagggcgaggagctgttcaccggggtggtgcccatccaggtcgagctggacggcgacgtaaacggccacaagttcagcgtgtccggcgagggtgagggcgatgccacctacggcaagctgaccctgaagttcatctgcaccaccggcaagctgcccgtgccctggcccaccctcgtgaccaccctgacctacggcgtgcagtgcttcagccgctaccccgaccacatgaagcagcacgacttcttcaagtccgccatgcccgaaggctacatccaggagcgcaccatcttcttcaaggacgacggcaactacaagacccgcgccgaggtgaagttcgagggcgacaccctggtgaaccgcatcgagctgaagggcatcgacttcaaggaggacggcaacatcctggggcacaagctggagtacaacactagtaagcttgtcgccacgaacttctctctgttaaagcaagcaggagacgtggaagaaaaccccggtcctatggtgtctaagggcgaagagctgattaaggagaacatgcacatgaagctgtacatggagggcaccgtgaacaaccaccacttcaagtgcacatccgagggcgaaggcaagccctacgagggcacccagaccatgagaatcaaggtggtcgagggcggccctctccccttcgccttcgacatcctggctaccagcttcatgtacggcagcaagaccttcatcaaccacacccagggcatccccgacttctttaagcagtccttccctgagggcttcacatgggagagatccaccacatacgaagacgggggcgtgctgaccgctacccaggacaccagcctccaggacggctgcctcatctacaacgtcaagatcagaggggtgaacttcccatccaacggccctgtgatgcagaagaaaacactcggctgggaggccagcaccgagatgctgtaccccgctgacggcggcctggaaggcagagactacatggccctgaagctcgtgggcgggggccacctgatctgcaacgccaagaccacatacagatccaagaaacccgctaagaacctcaagatgcccggcgtctactatgtggacagaagactggaaagaatcaaggaggccgacaaagagacctccgtcgagcagcacgaggtggctgtggccagatactgcgacctccctagcaaactggggcacaaactttaataaGAATTC.

pAAV-hSyn-DIO-mCherry was a gift from Bryan Roth (Addgene plasmid cat. no. 50459; http://n2t.net/addgene:50459; RRID:Addgene_50459).

### Perfusion and tissue preparation

After the completion of behavioral experiments or after 4 weeks post-surgery for histological experiments, the mice were deeply anesthetized using 5% isoflurane. Then, the mice were transcardially perfused with 4% paraformaldehyde (PFA; Biotrend Chemikalien, cat. no. 30400000-2) in phosphate-buffered saline (PBS). Mice were decapitated and the brain was dissected out of the skull. The brain was post-fixed in PFA for 24 h at 4 °C. Brains were sectioned coronally at 100 µm using a vibratome (Leica VT1000S). Sections were mounted directly using medium containing DAPI (Fisher Scientific, cat. no. 15395816) onto adhesion slides. Images were acquired using a Vectra Polaris imaging system (Akoya Biosciences).

### Injection site heatmaps for behavioral testing

Viral expression was plotted based on mCherry expression in coronal sections using a custom-written MATLAB script (MathWorks, version R2022a). The sections were aligned, background-subtracted, binarized and overlaid across animals for the main injection site. For each pixel, expression probability was calculated as the fraction of animals showing signal at that location, generating heatmaps scaled from 0 to 1.

### Anterograde tracing

For anterograde tracing, 30 nL of AAV-2/2-EF1a-DIO-hChR2(H134R)-mCherry-WPRE-hGHp(A) (3.1*10^12 gc/mL; UNC vector core) was injected into the TeA of *N* = 6 Adcyap1r1-Cre mice. 4 weeks after the surgery, mice were transcardially perfused with 4% PFA and the whole brain was embedded in 4% agarose and coronally sectioned into 50 µm slices as described previously.

For the quantitative assessment, the 18 brain areas exhibiting the highest density of axonal labeling were selected for analysis. For each region, the coronal section showing the densest fluorescent axonal signal was selected. Regions of interest (ROIs) were manually delineated in ImageJ by aligning the sample slice with the corresponding reference from the Allen Brain Institute atlas (Allen Institute for Brain Science, 2011). This alignment was performed using Affinity Designer.

Subsequently, each ROI was manually thresholded to discriminate the axonal signal from tissue autofluorescence. Area coverage was calculated as the percentage of positive pixels within each ROI. Furthermore, the total number of positive pixels was summed across all 18 regions per mouse to calculate the relative percentage of the axonal innervation across the brain in each area.

### RNA sequencing analysis

Publicly available single-cell RNA sequencing (scRNA-seq) datasets from Yao et al. (2021) were analyzed with custom-written MATLAB scripts to assess *Adcyap1r1* and *Adcyap1* expression across cortical brain areas and across cell types. Data were downloaded from GEO accession number GSE185862. We used original cell type assignments as provided in the metadata. Expression levels were averaged across all cells within each cell type. scRNA-seq data were analyzed as log(TPM+1).

### Fluorescent *in situ* hybridization

C57BL/6JRj mice were anesthetized using 5% isoflurane. The brains were quickly dissected and frozen in Tissue Tek OCT compound on dry ice. Subsequently, brains were sectioned into 20 µm thick coronal sections, adhered to SuperFrost Plus slides, and immediately refrozen. The samples were fixed in 4% PFA for 15 min at 4 °C. For protease treatment, slices were incubated in Protease III for 20 min. The samples were then hybridized at 40 °C with target probes: Mm-*Adcyap1r1*-C1 (biotechne, cat. no. 409561, of note, this probe is targeting a sequence present in all *Adcyap1r1* mRNA splice variants), Mm-*Sst* (biotechne, cat. no. 404631-C2), Mm-*Adcyap1*-C2 (biotechne, cat. no. 405911-C2), Mm-*Aqp4*-C2 (biotechne, cat. no. 417161-C2), Mm-*Pvalb* (biotechne, cat. no. 421931-C3), Mm-*Vip* (biotechne, cat. no. 415961-C3), Mm-*Slc17a7* (biotechne, cat. no. 416631-C3), Mm-*Gfap*-C3 (biotechne, cat. no. 313211-C3). To avoid bleed-through into other fluorescent channels, Mm-*Vip* and Mm-*Sst* probes were diluted 1:10 using probe dilutant (RNAScope). Mm-*Pvalb* was diluted 1:4. Fluorophore reagents were diluted to various concentrations (1:1500 and 1:30000) depending on color and probe. The following fluorophores were used: Opal 520 (Akoya, cat. no. FP1487001KT), Opal 570 (Akoya, cat. no. FP1488001KT), Opal 690 (Akoya, cat. no. FP1497001KT). Slides were coverslipped with ProLong antifade reagent (Fisher Scientific, cat. no. 11539306).

Images were taken at a Leica SP8 X confocal microscope using a 63x 1.4 NA oil immersion objective (Harvard NeuroDiscovery Center), at a pixel size of 180 nm and an optical section of 0.9 μm.

Manual cell quantification was completed using ImageJ. A cell was considered positive for *Adcyap1r1*, if at least 2 clearly visible mRNA puncta were present.

### Electrophysiological recordings and epifluorescence slice imaging

The mice were anesthetized using 5% isoflurane and transcardially perfused with 30 mL cooled sucrose solution (212 mM sucrose, 3 mM KCl, 1.25 mM NaH_2_PO_4_, 26 mM NaHCO_3_, 1 mM MgCl_2_, 0.2 mM CaCl_2_, 10 mM glucose) saturated with carbogen gas (95% O_2_ / 5% CO_2_). The brains were dissected and sectioned into 300 µm thick coronal slices in cold carbogenated sucrose solution using a vibratome (Leica VT1200). The brain slices were kept in carbogenated ACSF (Artificial cerebrospinal fluid; 125 mM NaCl, 25 mM NaHCO_3_, 1.25 mM NaH_2_PO_4_, 2.5 mM KCl, 2 mM CaCl_2_, 1 mM MgCl_2_, 25 mM glucose) for 10 min at 37 °C and subsequently maintained at room temperature until use.

Recordings were performed using an Olympus BX51WI microscope equipped with an ORCA-Fusion Digital CMOS camera (Hamamatsu, cat. no. C14440-20UP). A 10 x water-immersion objective (Olympus, UMPLFLN10XW) was used for slice imaging and a 40 x objective (Olympus, LUMPLFLN40XW) for electrophysiological recordings. During recordings, slices were continuously perfused with carbogenated ACSF at ∼32 °C.

#### Whole-cell patch-clamp recordings

Whole-cell patch-clamp recordings were performed with pipettes pulled from borosilicate glass capillaries with resistances of 2-5 MΩ. Cells were visualized with an upright microscope equipped with differential interference contrast (DIC) and a pT-100-770 LED (cooled, cat. no. pT-100-770-50E-20) for IR illumination. All electrophysiological recordings were acquired using an EPC 10 USB Double Patch Clamp Amplifier. Signals were sampled at 10 kHz using Patchmaster Next software. Liquid junction potentials were not corrected for. Pipettes and microscope movements were controlled with Luigs&Neumann instruments. Pyramidal cells were identified by their cell shape, localization, input resistance, membrane capacitance and firing pattern. Fast firing patterns were acquired for identification of cell types using 1 s long current pulses with 2 s intersweep interval, starting at -250 pA and incrementally increasing by 150 pA until saturation. Postsynaptic currents (PSCs) were recorded in identified cells voltage-clamped at −70 mV. For inhibitory PSCs (IPSCs), NBQX (10 μM, TargetMol cat. no. T4113-200 mg) and R-CPP (10 μM, Tocris cat. no. 0247/50) were added to the bath, and K^+^-based, high Cl^-^intracellular solution was used (in mM: 127.5 KCl, 11 EGTA, 10 HEPES, 1 CaCl2, 2 MgCl2, 2 Mg-ATP and 0.3 GTP, pH 7.3 adjusted with KOH). For excitatory PSCs (EPSCs), identified cells were voltage-clamped at −70 mV and gabazine (10 μM, SR 95531 hydrobromide, Tocris cat. no. 1262/50) and CGP 55845 (10 μM, Tocris cat. no. 1248/50) were added to the bath. K^+^-based, low-Cl^-^intracellular solution was used (in mM: 130 K-Gluconate, 2 EGTA, 10 HEPES, 4 Mg-ATP, 0.3 GTP-Na, 10 Phosphocreatine-Na, 10 Na-Gluconate, 4 NaCl, pH 7.2 adjusted with KOH). For optogenetic stimulation experiments in acute slices, AAV 2/2-EF1a-DIO-hChR2(H134R)-mCherry-WPRE-hGHp(A) was stereotaxically injected. Putative target cells of ChR2-labeled neurons were patched in close proximity to labeled axons (<20 μm). 473 nm light (Cobolt 473 nm MLD, Cobolt/Huebner) at maximally 31 mW was delivered through a 40 x objective. 0.1-5 ms lasting pulses were used for local connectivity with 50 ms inter pulse interval and 10 s inter trial interval. Light power for each experiment was adjusted to trigger PSCs reliably and with a clear monosynaptic peak. 5 ms pulses and 31 mW were used for long-range connectivity. Amplitudes of PSCs were defined as the negative peak within an approximately 15 ms time window from the start of the light pulse. Paired pulse ratio was defined as the amplitude to the second light pulse divided by the amplitude to the first light pulse after applying a sliding average to the amplitudes with a window size of 1 min. Amplitudes and frequencies were normalized to the first 3 minutes of the recording. Spontaneous EPSCs (sEPSCs) were recorded with K^+^-based, low-Cl^-^intracellular solution (see above) without any additional antagonists in the ACSF. Neurons were held at -70 mV continuously except during series resistance measurement that happened every 5 s. The 105 ms time window for the series resistance measurement was cut out from the trace for analysis of sEPSCs. sEPSCs were detected with a custom-made MATLAB script that detects negative peaks based on noise level, peak amplitude and duration of the transients surrounding the peaks. Amplitude was defined as the difference between negative peak and maximum value within a 10 ms time window before the negative peak, and by subtracting the defined noise level. Amplitudes and frequency were smoothed across 20 s time windows. Frequency was then normalized to the first 3 min of the recording (baseline). Voltage ramp recordings were used to measure currents at different potentials. To this end, neurons were held between -140 and -45mV with a voltage change rate of 1 mV/ms. For visualization, a range of -130 to -50 mV was chosen, to avoid contamination with capacitive currents at the initial voltage step, and to avoid contamination with activation of voltage-dependent sodium channels at action potential threshold. A current ramp (current increased at a rate of 100 pA per second) was used to determine the rheobase. The rheobase was defined as the current needed to evoke the first action potential. The duration of the ramp (and thus the final current) was adjusted to each neuron’s baseline rheobase. Resting membrane potential was defined as the mean voltage at 0 pA holding potential across a 500 ms time period, measured every 30 s. Rheobase and resting membrane potential and voltage ramp responses were averaged across 3 min starting 8.5 min after the start of the recording. K^+^-based low-Cl^-^solution (see above) was used. 300 nM PACAP (PACAP-38, Phoenix Pharmaceuticals; cat. no. 052-05) were bath applied for 2 min (from minute 3 to 5) unless indicated otherwise. Where indicated, 10 µM of Forskolin (CliniSciences, cat. no. B1421-100mg) was bath applied from minute 3 to 5. Note that we did not correct for the 1 min delay for the peptide to reach the slice chamber. The following drugs were added to the ACSF to identify PACAP-evoked signaling mechanisms: ZD7288 (20 μM, TargetMol cat. no. T7516-25 mg), riluzole hydrochloride (10 μM, Hellobio cat. no. HB0548-10mg), U0126 (20 μM, TargetMol cat. no. T21332-25 mg), H-89 dihydrochloride (10 μM, THP Medical Products cat. no. MCE-HY-15979A-10mg or Tocris, cat. no. 2910). Series resistance was continuously monitored in voltage-clamp mode measuring peak currents in response to small hyperpolarizing pulses. Recordings with series resistance changes of more than 25% were discarded. Series resistances of 35 MΩ were accepted for analyzing PSCs in interneurons and 25 MΩ were accepted for pyramidal cells. MATLAB was used for offline analysis of all data.

#### Slice imaging

For slice imaging, genetically encoded fluorescent sensors (see above) were used. A GFP (green fluorescent protein) filter set was used to control the emission and excitation spectra (AFH, cat. no. F46-002; excitation 470/40 nm, emission 525/50 nm, 495 nm beam splitter T495 LPXR; AFH, cat. no. F48-495) of green fluorescent protein (GFP)-based reporters. Red fluorescent proteins (RFP) were imaged using an RFP filter set (excitation 575/32 nm, emission 625/32 nm, 604 nm beam splitter; AHF). Micromanager software (D. Edelstein et al., 2014) was used to synchronize camera shutter and the excitation LED. Images were acquired at a framerate of 4 Hz with 50-100 ms exposure time per frame.

The first 50 frames of each recording were acquired in the RFP channel to document reporter-independent expression where applicable.

An area with clear cellular fluorescence within the TeA was selected for imaging and pre-recording of 5-10 min was performed, allowing the slices to stabilize in the conditions and reduce steep bleaching curves. For recordings performed in astrocytes, ACSF contained tetrodotoxin (TTX; 500 nM; Tocris, cat. no. 1069/1) to block action potential-dependent neuronal activity. For recordings performed in neurons, NBQX (10 µM, TargetMol, cat. no. T4113-200 mg), CPP (10 µM, Hellobio, cat. no. HB0021), CGP 55845 (10 µM, Hellobio, cat. no. HB0960), and gabazine (10 µM, Hellobio, cat. no. HB0901) were added to the bath to prevent recruitment of neuronal circuits that could lead to endogenous PACAP release. For experiments probing intracellular signaling in response to PACAP, slices were continuously exposed to specific pathway inhibitors: The PI-phospholipase-C (PLC) blocker U73122 (5 µM; Tocris, cat. no. 1268/10), the PC-PLC blocker D609 (10 µM; Tocris, cat. no. 1437/10), the Gq inhibitor YM-254890 (500 nM; Tocris cat. no. 7352), the IP_3_R inhibitor 2-APB (100 µM; Tocris, cat. no. 1224/10), the protein kinase A (PKA) inhibitor H-89 dihydrochloride (10 µM; THP Medical Products cat. no. MCE-HY-15979A-10mg or Tocris, cat. no. 2910), the L-type channel Ca^2+^ inhibitor nimodipine (10 µM; Tocris, cat. no. 0600), the phosphodiesterase inhibitor IBMX (1 mM; Sigma Aldrich, cat. no. 410957-250MG) and the Gβ/γ inhibitor gallein (20 µM; Tocris, cat. no. 3090).

The experimental recording was 27-28 min long. For most experiments, PACAP-38 (30–3000 nM; Phoenix Pharmaceuticals; cat. no 052-05, dissolved in ddH2O) was infused from minute 3-5. Note that PACAP-38 was chosen over the alternative peptide form PACAP-27 due to its more ubiquitous potency (Spengler et al., 1993). Serving as controls, all experiments were repeated with infusing a matching volume of vehicle (ddH2O, 0 nM PACAP) to exclude changes in fluorescence due to valve switching and related pressure differences. For experiments testing AAV-hGFAP-chI-bBARK(rgs)-P2A-mCherry or its control, norepinephrine (10 µM; TargetMol, cat. no. TGM-T0871-50mg) was bath-applied. To test whether PAC1 was activated, 150 nM of the specific PAC1 agonist Maxadilan (Bachem; cat. no. 4067317) was bath applied. From minute 15 to the end of the recording, 10 µM of Forskolin for cAMP imaging (CliniSciences, cat. no. B1421-100mg) or 30 mM KCl (Fisher Scientific, cat. no. 10735874) for calcium imaging were added as positive controls to confirm slice viability.

#### Slice imaging data analysis

Data analysis was performed using custom-written MATLAB scripts (MathWorks, version R2022a). Fluorescence signals were extracted from manually defined regions of interest (ROIs) corresponding to individual fluorescent neuronal cells or astrocyte “domains” (including cell bodies and processes) within the TeA. Fluorescence changes were expressed as ΔF/F by normalizing fluorescence changes to the mean baseline fluorescence (F_0_) during the first 3 min of recording for all experiments.

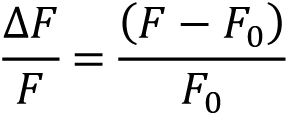

Because cADDis is an inverse-response cAMP sensor, in which an increase in intracellular cAMP produces a decrease in fluorescence, cADDis responses were sign-inverted for visual representations such that increases in inferred cAMP were represented as positive-going signals.

To correct for slow fluorescence bleaching during imaging, an exponential decay function was fitted to the lower 25% percentile of the vehicle-control (ddH2O) traces and subtracted from all recordings for astrocytic recordings, thereby removing gradual baseline drift while preserving stimulus-evoked responses. For neurons, a bleaching curve was estimated for each recording by fitting a single-exponential decay function to the mean normalized fluorescence trace across ROIs.

Slices that failed to respond to 30 mM KCl or 10 µM Forskolin application were excluded from further analysis. For quantification of ligand-evoked responses, fluorescence traces were averaged across all ROIs within a slice to obtain a single representative trace per slice. To reduce the influence of spontaneous fluctuations on peak responses, each slice trace was smoothed using a moving-average filter over a 1.2 s moving window. Response amplitude was then defined as the maximum smoothed ΔF/F value occurring within 4-15 min of the recording. Responses were compared against control recordings in which matched volumes of ddH2O were bath-applied. For Ca^2+^ imaging recordings in neurons, peak ΔF/F values during baseline (0-3 min) and response (4-15 min) were extracted for each recording and statistically compared. For statistical comparisons, ≥ 6 slices from ≥ 3 animals per condition were collected. Data are reported as mean ± SEM across slices unless stated otherwise.

Time to half-maximum was calculated as the first time point at which the response reached 50% of the amplitude between the baseline at the point when PACAP reaches the recording chamber and the maximum response observed prior to Forskolin application (between 4 and 15 min).

### Auditory fear conditioning

Female mice in the age of 8-12 weeks underwent stereotaxic AAV injections as described above.

Prior to experimental testing, mice were handled for 7 days to get habituated to the experimenter. On day 2 of handling, mice were single-housed.

Behavioral experiments were conducted between Zeitgeber time (ZT) 1 and 5 in custom-built conditioning boxes consisting of a shock-grid floor enclosed by acrylic plates (31 x 23.5 x 37 cm; length x width x height). Each box was equipped with a Blackfly S USB3 camera (BFS-U3-13Y3M-C, Teledyne), visible and infrared light sources, and a speaker. Auditory stimuli were played from a Titanium Dome Tweeter (TW025A20 1, Audax) driven by a SparkFun MP3 player shield (12660. SparkFun Electronics) and calibrated to 55 dBA at mouse height. Stimulus presentation, lights and shock delivery were controlled via an Arduino microcontroller and logged using a custom MATLAB script through a T7 DAQ Board (LabJack). Videos were recorded at 30 frames per second.

The fear conditioning paradigm was adapted from Letzkus et al. (2011) and spanned three consecutive days. All sessions were conducted during the early light phase. On day 1 (habituation), mice explored the chamber for 10 min without stimuli. On day 2 (conditioning), a 3 min baseline was recorded, followed by presentation of two complex sounds (upsweeps or downsweeps; 15 repetitions each; 500 ms sweeps ranging from 5-10 kHz with 45-180 s interstimulus intervals). 15 repetitions of each sound were delivered in pseudorandom order. One sound (CS^+^) co-terminated with a 1 s foot shock (0.6 mA), whereas the other (CS^-^) remained unpaired. Assignment of upsweeps versus downsweeps as CS^+^ or CS^-^was counterbalanced across animals. On day 3 (retrieval), mice were weighed and injected intraperitoneally (i.p.) with CNO (4 mg/kg in 0.9% NaCl solution (saline); HelloBio, cat. no. HB6149-50mg) or saline (control) 30 min prior to testing for the chemogenetic experiment. Contextual cues were altered to reduce contextual fear: the grid floor was covered with white plastic, the chamber was reshaped into a semicircle, the visible light was turned off, and a vanilla odor was introduced.

After a 3 min baseline, mice received 15 CS^+^ and 15 CS^-^presentations in the same order as during conditioning, without shock delivery.

#### Analysis

A custom-written MATLAB script was used to quantify freezing levels on the retrieval day. Freezing was defined as no movement of the mouse except for breathing for ≥ 2 s. Briefly, a binary mask of the mouse was generated and frame-to-frame pixel changes were calculated using centroid-based tracking. Thresholds distinguishing freezing from movement were semi-automatically determined for each video based on presentation of predicted freezing segments with an increasing or decreasing number of pixel changes. Ambiguous freezing segments close to defined thresholds were semi-automatically validated. Freezing in response to CS^+^ and CS^-^was calculated by averaging freezing levels across all sound presentations (15 trials per stimulus). Freezing values reflect absolute freezing levels and were not baseline-subtracted. Baseline freezing was quantified during the 3 min period preceding stimulus presentation.

#### Photometry

Mice implanted with fiber-optic cannulae experienced the same behavioral paradigm as described above. During the final two days of handling, mice were habituated to gentle manipulation of the implant and patch cord connection.

Immediately before behavioral testing, implanted fiber-optic cannulae were connected to a 200 µm diameter, 5.5 m long patch cord (NA 0.37; Doric Lenses, MFP_200/220/900–0.37_5.5m_FCM-MF1.25, low autofluorescence epoxy). Excitation light was provided by a blue LED (470 nm; fiber-coupled LED; Thorlabs; cat. no. M470F3). The excitation light path was guided through a Fluorescence MiniCube (Doric lenses; iFMC5-G2_E1(460-490)_F1(500-540)_E2(555-570)_F2(580-680)_S). Excitation light power at the patch cord tip was set to 45 µW. Signals were detected and amplified using Doric’s built-in photodetectors and amplifiers, acquired using a T7 Daq Board (LabJack), and recorded with custom MATLAB software. LED driver output was sinusoidally modulated at 171 Hz, and data were sampled at 2052 Hz as described previously by Melzer et al. (2021) and Owen & Kreitzer (2019).

Fluorescence signals were demodulated by extracting the spectral power at the respective modulation frequencies from a short-time Fourier transform computed in MATLAB using a 200-sample sliding window and 180-sample overlap. To correct for background autofluorescence originating from the optical setup and patch cord, fluorescence power was measured prior to connecting the patch cord to the implanted fiber and subtracted from all recorded signals before further analysis. Photobleaching was corrected by fitting an exponential decay to the minima of sliding 6.8 s windows. The trace was then divided by the fit. ΔF/F was computed relative to baseline fluorescence calculated as the mean of the first 2 min of recording prior to sound presentation.

For sound-aligned analyses, fluorescence traces were segmented into trials time-locked to CS onset and averaged within each mouse. For shock-aligned analyses, traces were aligned to foot shock onset during conditioning. For freezing-aligned analyses, ΔF/F traces were aligned to freezing onset and offset, defined from the freezing annotation.

Peak response amplitude was defined as the maximum ΔF/F value within a predefined post-stimulus window (0-29 s after CS onset for sound-evoked responses; 0-10 s after shock onset for shock-evoked responses). Peak amplitudes were first computed for each mouse based on the trial-averaged trace and then averaged across mice.

For testing attenuation or amplification to sounds, sound-evoked responses were averaged within each mouse across early (trials 1-5), middle (trials 6-10) and late (trials 11-15) trial blocks. Peak amplitudes were extracted from the trial-averaged trace for each block, resulting in one value per mouse and condition and were compared using one-way repeated-measures ANOVA.

To test whether shock neuronal responses were correlated with shock-induced locomotion, maximum ΔF/F and maximum speed within 0-10 s after shock were correlated, and linear regression, Pearson correlation and cross-correlation analyses were performed using custom-written MATLAB scripts. For speed calculation, the centroid location of each mouse was computed for each frame, and the frame-to-frame distances calculated with Euclidean distance formula. After applying a sliding average with a window size of 2 sample points, distances were converted into cm/s units.

To determine correlation between fluorescence CS^+^ responses and freezing during retrieval, mean responses during CS^+^ and mean freezing during CS^+^ were averaged per mouse across all sounds during retrieval. To test whether fluorescence responses to CS^+^ during conditioning can predict freezing levels during retrieval, peak responses during fear conditioning CS^+^ (excluding the last second coinciding with the shock) and mean freezing levels during CS^+^ during retrieval were averaged across all 15 CS^+^ per mouse.

#### Exclusion criteria

Mice were excluded when histology revealed a lack of viral expression or a misplaced injection site, with the majority of AAV expression on either hemisphere visible outside of the TeA, or showing spread into non-cortical adjacent regions (e.g., hippocampal formation). Other exclusion criteria were technical failures during the behavioral experiment or insufficient fluorescence (baseline fluorescence <3x autofluorescence for fiber-implanted mice).

### Quantification and statistical analysis

Group sizes are indicated in figure legends; power calculations were used to predetermine sample sizes.

All analyses were performed in MATLAB using custom-written scripts unless indicated otherwise. Data are presented as mean ± SEM or median [IQR]. All statistical tests were two-sided. Normality was assessed using Shapiro-Wilk tests and homogeneity of variance using *F*-tests. For comparisons between two independent groups, unpaired Student’s *t*-tests were used when data were normally distributed and variances were homogeneous. When data were normally distributed but variances differed between groups, Welch’s unpaired *t*-test was used. When assumptions for parametric testing were not met, Mann-Whitney *U* tests or Kruskal-Wallis tests were used, as appropriate. Where multiple related hypotheses were tested, *p*-values were adjusted for multiple comparisons using the Bonferroni-Holm method with family-wise α = 0.05. For statistical comparisons of freezing data, *p*-values were corrected for testing of three related measures: baseline freezing, CS^+^ freezing, and CS^-^freezing.

For auditory fear conditioning experiments, behavioral readouts were analyzed on a per-mouse basis and compared between experimental and control groups.

#### Blinding strategy

For all behavioral experiments, experimenters were blinded to group assignment throughout the entire experimental workflow, including surgeries, behavioral testing, analysis, histological validation and exclusion of mice. Mice were randomly assigned to groups. Groups that were statistically compared in this study were run during the same time period and, wherever possible, in parallel.

## Data availability

Data reported in this paper will be shared by the lead contact upon request. Any additional information required to reanalyze the data reported in this paper is available from the lead contact upon request.

## Code availability

All original code is available from the lead contact upon request.

## Acknowledgements

We thank Philipp Velicky and the team of the Imaging Core Facility for technical support. We thank Bryan Roth, Bernardo Sabatini, Karl Deisseroth and Baljit Khakh for providing plasmids. We thank the animal care staff (Sonja Reynoso de Leon, Aurica Jelinek, Corina Brottrager, Christian Schönauer and Jutta Pilecky) for their support.

This work was funded by the Vienna Science and Technology Fund (WWTF) and the City of Vienna through project VRG21-015.

## Author contributions

Conceptualization: S.N.H., S.M., Data analysis: S.N.H., S.M., A.C.M., B.D., V.B., Investigation: S.N.H., S.M., A.C.M., B.D., V.B, Writing: S.N.H., S.M, Visualization: S.N.H., S.M., A.C.M, Supervision: S.M., Funding acquisition: S.M.

## Ethics declaration

Competing interests: None to be declared.

## Extended Data

**Extended Data Fig. 1.**
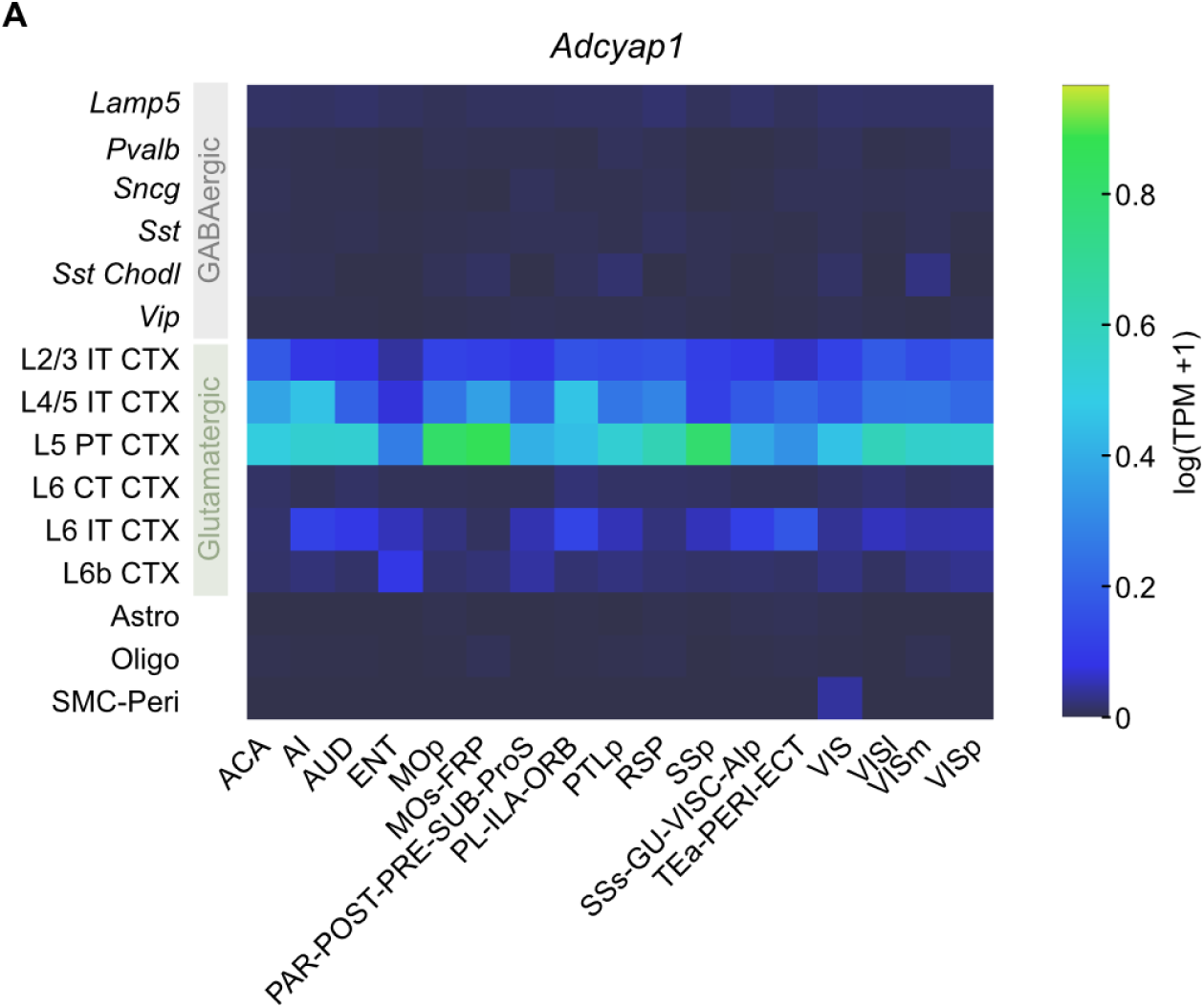
**A.** Heatmap showing expression (log(TPM+1)) of *Adcyap1* across major GABAergic, glutamatergic and non-neuronal cell types in 17 cortical area groups. Expression data and cell-type classification were extracted and reanalyzed from Yao et al 2021. Brain area abbreviations as in Fig. 1.

**Extended Data Fig. 2.**
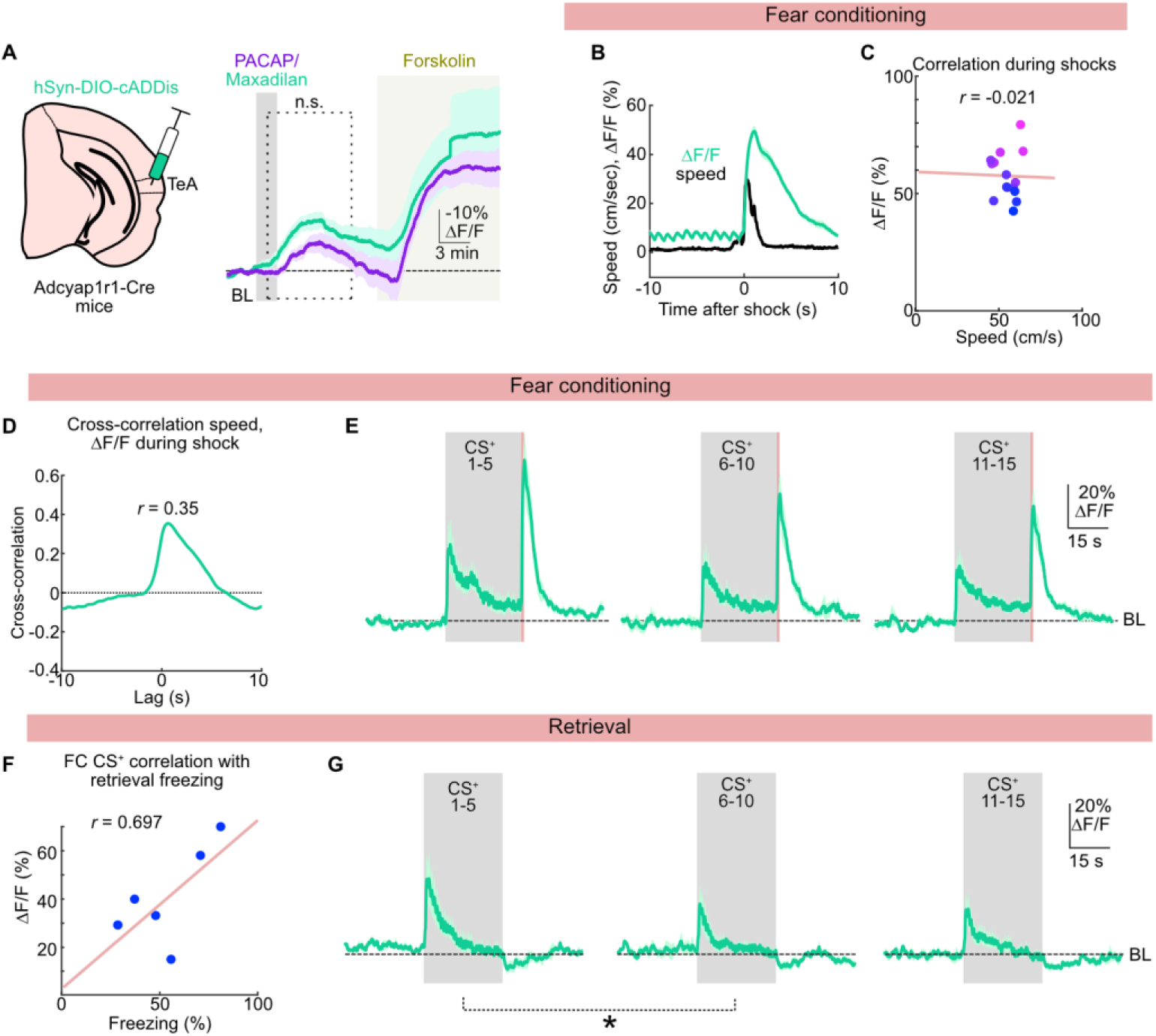
**A.** Left: Schematic of AAV-injections of AAV-hSyn-DIO-cADDis into TeA in Adcyap1r1-Cre mice. Right: Averaged fluorescence cADDis signals (-ΔF/F) upon PACAP (300 nM) and maxadilan (150 nM) application. Peak signals: PACAP -34.11 ± 6.7% ΔF/F; maxadilan: -62.95 ± 17.85% ΔF/F, *t*(6.4) = 1.51, *p* = 0.356; unpaired *t*-test, *N* = 6 -7 slices per condition. **B.** Mean speed and ΔF/F during the shock period (±10 s), averaged across mice, *N* = 6 mice. **C.** Correlation between maximum locomotor speed and maximum Ca²⁺ transients across all shocks (+10 s), averaged across mice. Each point represents the mean response to a given shock (first shock (pink) to 15^th^ shock (blue)). Linear regression and Pearson correlation coefficient are shown, *N* = 6 mice. **D.** Cross-correlation between locomotor speed and ΔF/F during the shock period (±10 s), averaged across mice, *N* = 6 mice. **E.** Normalized fluorescence changes (ΔF/F) from *Adcyap1r1*-Cre^+^ neurons expressing GCaMP6s aligned to CS⁺ presentations during fear conditioning. Traces are grouped by trial number (CS⁺ 1–5, 6–10, and 11–15). Peak ΔF/F CS⁺ 1-5: 38.51 ± 11.74%, peak ΔF/F CS^+^ 6–10: 29.68 ± 8.15%; peak ΔF/F CS^+^ 11-15: 27.48 ± 7.08%. No statistical difference in peak ΔF/F during CS^+^ presentations across trial blocks was detected (*F*(2,8) = 3.015; *p* =0.094; one-way repeated-measures ANOVA), *N* = 6 mice. **F.** Quantification of correlation between average Ca^2+^ transients during CS^+^ on conditioning day and freezing level during retrieval session in *Adcyap1r1*-Cre^+^ neurons. Dot plot with linear regression and Pearson correlation, *N* = 6 mice. **G.** Normalized fluorescence changes (ΔF/F) in *Adcyap1r1*-Cre^+^ neurons expressing GCaMP6s aligned to CS⁺ presentations during retrieval, grouped by trial number (CS⁺ 1-5, 6-10, and 11-15). Peak ΔF/F CS⁺ 1-5: 36.02 ± 10.52%; peak ΔF/F CS⁺ 6-10: 23.94 ± 8.17%; peak ΔF/F CS⁺ 11-15: 25.21 ± 6.46%. A statistical difference in peak ΔF/F during CS^+^ presentations across trial blocks was detected (*F*(2,8) = 5.558, *p* = 0.024; one-way repeated-measures ANOVA). Bonferroni-Holm corrected post-hoc paired *t*-tests revealed a significant difference comparing peak ΔF/F CS⁺ 1-5 with CS⁺ 6-10 (*t*(5) = 3.829; *p* = 0.037), *N* = 6 mice. Data are shown as mean ± SEM.

**Extended Data Fig. 3.**
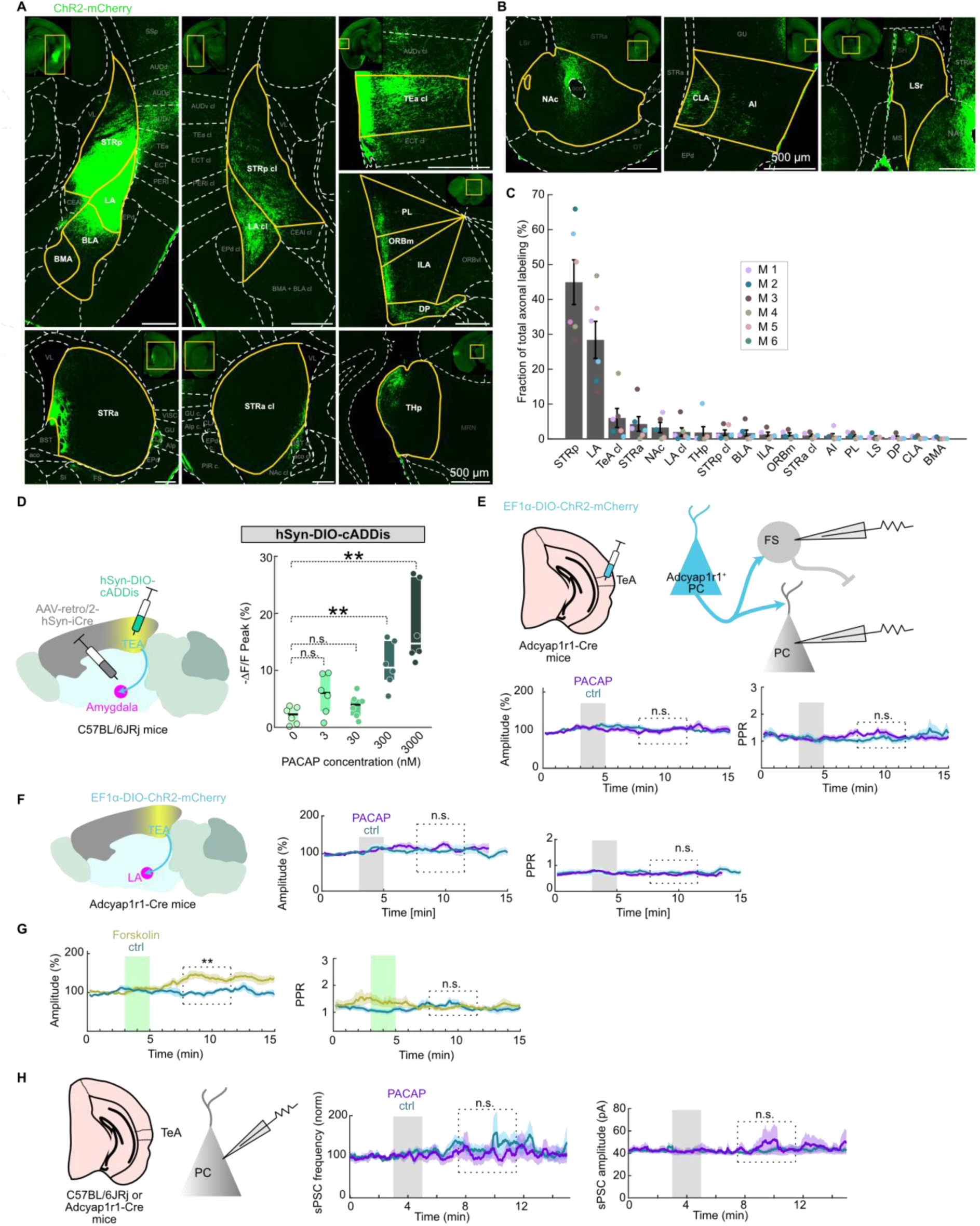
**A.-B.** Exemplary epifluorescent images of axonal labeling in target areas, *N* = 6 mice. **C.** Quantification of axonal labeling area as percent of total labeling across all quantified areas. Same color code as in Fig. 3C. **D.** Schematic and averaged peak fluorescence cADDis signals (-ΔF/F) of TeA-amygdala projection neurons upon different concentrations of PACAP application. Statistical comparison with 0 nM / ctrl.0 nM / ctrl: -2.1 ± 0.622% ΔF/F; 3 nM: -5.77 ± 1.44% ΔF/F, *t*(10) = -2.344, *p* = 0.082; 30 nM: -3.79 ± 0.86% ΔF/F, *t*(10) = -1.599, *p* = 0.141; 300 nM: -10.96 ± 1.62% ΔF/F, *t*(10) = -5.097, *p* = 0.0014; 3000 nM: -17.85 ± 2.85% ΔF/F, *t*(10) = -5.405, *p* = 0.0012. Unpaired *t*-tests, *N* = 6 slices per condition. **E.** Top: Schematic of virus injection for patch-clamp experiments in principal cells (PC) and fast spiking neurons (FS). Bottom: Time course of amplitudes of optogenetically evoked EPSCs (oEPSCs, left) and paired pulse ratio (PPR, right) at glutamatergic synapses in the TeA. Gabazine and CGP were added to the bath to block GABAergic neurotransmission. Purple boxes indicate PACAP or vehicle (ctrl) application. Statistically compared time period indicated by dashed boxes. PACAP did not significantly change oEPSC amplitude (*t*(18) = -0.47, *p* = 0.64 (unpaired *t*-test)) or PPR (*t*(18) = 1.55, *p* = 0.14 (unpaired *t*-test)) at glutamatergic synapses: *N* = 10 cells per group. **F.** Left: Schematic of TeA-LA projection tested for presynaptic PACAP effects. Right: Average oEPSC amplitude and paired pulse ratio recorded from LA neurons receiving TeA input. Time period of PACAP or vehicle (ctrl) application indicated by grey bars. Statistically compared time period indicated by dashed boxes. No significant differences were detected for oEPSC amplitude (*t*(18) = -0.39, *p* = 0.70 (unpaired *t*-test)) or PPR (*t*(13) = 0.24, *p* = 0.82 (Welch’s unpaired *t*-test)). *N* =10 cells each. **G.** Positive control for experimental setup. Same as in **F.**, but using application of forskolin instead of PACAP to activate adenylate cyclase directly. Forskolin increased oEPSC amplitude significantly: *t*(14) = -3.66, *p* = 0.003 (unpaired *t*-test); PPR: *t*(14) = 0.47, *p* = 0.65. *N* = 8 cells per group. **H.** Left: Schematic of patch-clamp experiment. Right: Time course of spontaneous postsynaptic current (sPSC) frequency and amplitude in TeA upon PACAP or vehicle (ctrl) application (purple boxes). No additional drugs were added to the bath. *N* = 15 (ctrl), 12 (PACAP) cells. Statistics for frequency: *t*(25) = 0.59, *p* = 0.56 (unpaired *t*-test); amplitude: *t*(25) = -0.42, *p* = 0.68 (unpaired *t*-test). **Abbreviations**: ACO, anterior commissure; AI, agranular insular area; AUDd, dorsal auditory area; AUDp, posterior auditory area, AUDv, ventral auditory area; BLA, basolateral amygdala; cl, contralateral; BMA, basomedial amygdalar nucleus; BST, bed nuclei of the stria terminalis; CeA, central amygdala; cl, contralateral; CLA, claustrum; DP, dorsal peduncular area; ECT, ectorhinal area; EPd, endoprifirorm nucleus; GU, gustatory areas; ILA, infralimbic area; LA, lateral amygdala; LS, lateral septal nucleus; MG, medial geniculate complex; MRN, midbrain reticular nucleus; MS, medial septum; NAc, nucleus accumbens; ORBm, orbital area, medial part; ORBvl, orbital area, ventrolateral part; OT, olfactory tubercle; PAG, periaqueductul gray; PER, perirhinal area; PL, prelimbic area; SI, substantia innominata; STRa, anterior striatum; STRp, posterior striatum; SSp, primary somatosensory area; TeA, temporal association area; THp, posterior thalamus; VISC, visceral area; VL, lateral ventricle. All data shown as mean ± SEM.

**Extended Data Fig. 4.**
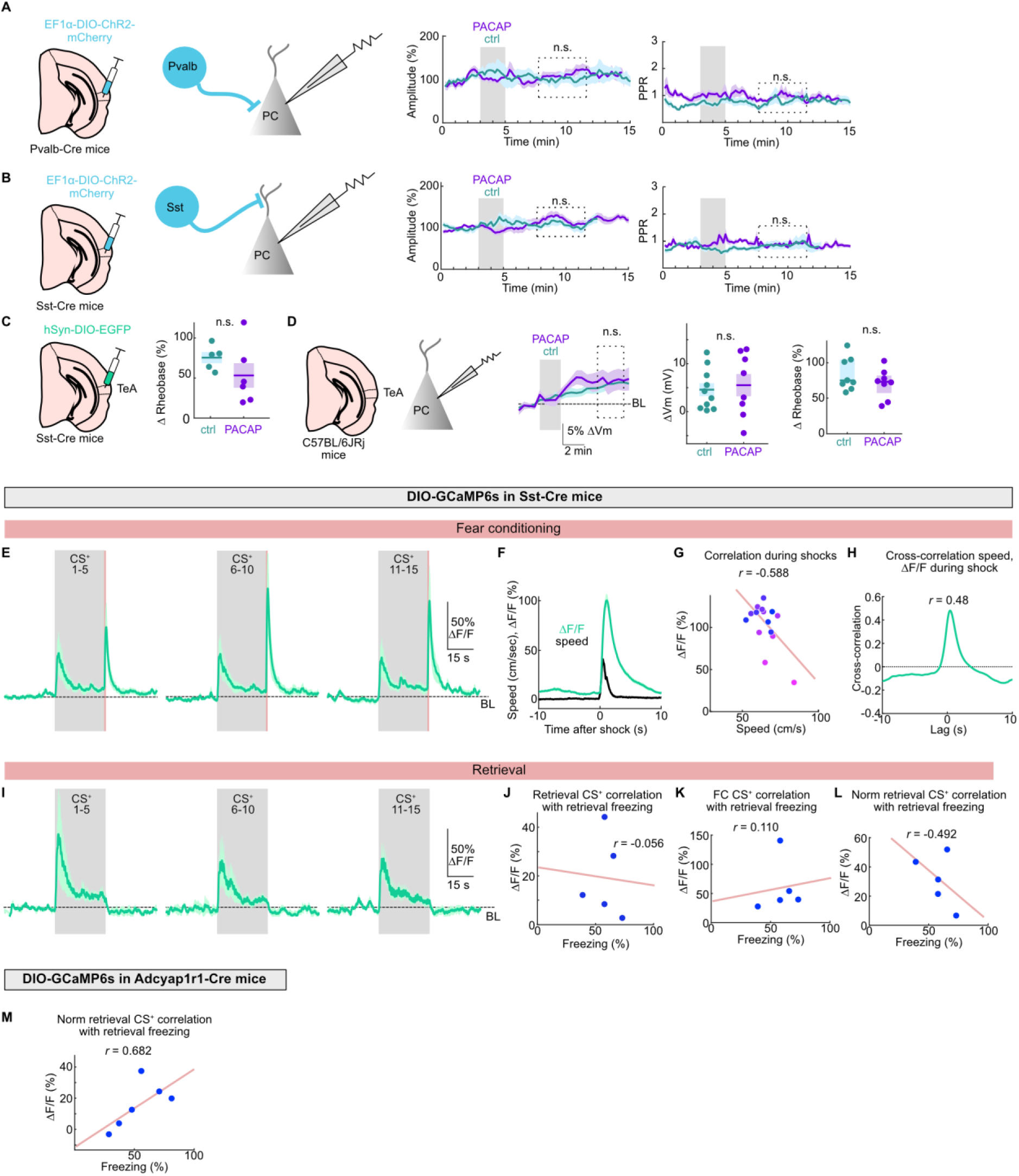
**A.** Left: Schematic of the viral injections and cell types targeted for optogenetic activation and electrophysiological recordings of inhibitory postsynaptic currents (IPSCs) in principal cells (PC). Right: Time course of amplitude and paired pulse ratio (PPR) of inhibitory postsynaptic currents. Application of PACAP and vehicle (ctrl) indicated by grey bars. Dashed box indicates analyzed period. No significant changes were detected for amplitude (*t*(16) = -0.76, *p* = 0.46, unpaired *t*-test) or PPR (t(16) = -0.86, *p* = 0.41, unpaired *t*-test). *N* = 9 cells each. **B.** Same as **A.**, but for optogenetic activation of Sst^+^ neurons. No significant changes were detected for amplitude (*U* = 25, *p* = 0.19, Mann-Whitney *U* test) or PPR (*U* = 35, *p* = 0.66, Mann-Whitney *U* test). *N* = 9 cells each. **C.** Schematic of whole-cell recordings from Sst^+^ neurons used to assess PACAP-induced changes in neuronal excitability (rheobase). Quantification of rheobase changes shown on the right. *t*(9) = 1.23, *p* = 0.25, unpaired *t*-test. *N* = 5 (ctrl) and 6 (PACAP) cells. **D.** Schematic of whole-cell recordings from principal cells (PC) used to assess PACAP-induced changes in neuronal resting membrane potential (middle) and excitability (rheobase, right). BL = mean baseline value. No significant changes were detected for membrane potential (*t*(16) = -0.36, *p* = 0.72, unpaired *t*-test) or rheobase (*U* = 28, *p* = 0.31, Mann-Whitney *U* test). *N* = 10 (ctrl) and 8 (PACAP) cells. **E.** Normalized fluorescence changes (ΔF/F) from Sst^+^ neurons expressing AAV-CBA-DIO-GCaMP6s aligned to CS⁺ presentations during fear conditioning. Peak ΔF/F CS⁺ 1-5: 51.49 ± 23.85%, peak ΔF/F CS^+^ 6–10: 55.94 ± 22.9%; peak ΔF/F CS^+^ 11-15: 55.13 ± 22.03%. No statistical difference in peak ΔF/F during CS^+^ presentations across trial blocks was detected (*F*(2,6) = 0.314; *p* = 0.739; one-way repeated-measures ANOVA), *N* = 5 mice. **F.** Mean speed and ΔF/F during the shock period (±10 s), averaged across the same mice as in E., *N* = 5 mice. **G.** Correlation between maximum locomotor speed and maximum Ca²⁺ transients across all shocks (+10 s), averaged across mice. Each point represents the mean response to a given shock (first shock (pink) to 15^th^ shock (blue)). Linear regression and Pearson correlation are shown, *N* = 5 mice. **H.** Cross-correlation between locomotor speed and Ca²⁺ ΔF/F during the shock period (±10 s), averaged across mice, *N* = 5 mice. **I.** Normalized fluorescence changes (ΔF/F) from Sst^+^ neurons expressing AAV-CBA-DIO-GCaMP6s aligned to CS⁺ presentations during retrieval. Peak ΔF/F CS⁺ 1-5: 87.34 ± 47.67%; peak ΔF/F CS⁺ 6-10: 57.75± 28.17%; peak ΔF/F CS⁺ 11-15: 58.98 ± 18.84%. No statistical difference in peak ΔF/F during CS^+^ presentations across trial blocks was detected (*F*(2,6) = 0.92; *p* = 0.405; one-way repeated-measures ANOVA), *N* = 5 mice. **J.** Quantification of correlation between average Ca^2+^ transients during retrieval CS^+^ and freezing level in Sst^+^ neurons. Dot plot with linear regression and Pearson correlation, *N* = 5 mice. **K.** Quantification of correlation between average Ca^2+^ transients during CS^+^ on conditioning day and freezing level during retrieval session in Sst^+^ neurons. Dot plot with linear regression and Pearson correlation, *N* = 5 mice. **L.** Same as **J.**, but CS^+^ responses during retrieval were normalized to mean CS^+^ responses during conditioning. **M.** Same as **L.**, but for *Adcyap1r1*-Cre^+^ neurons. Data are shown as mean ± SEM.

**Extended Data Fig. 5.**
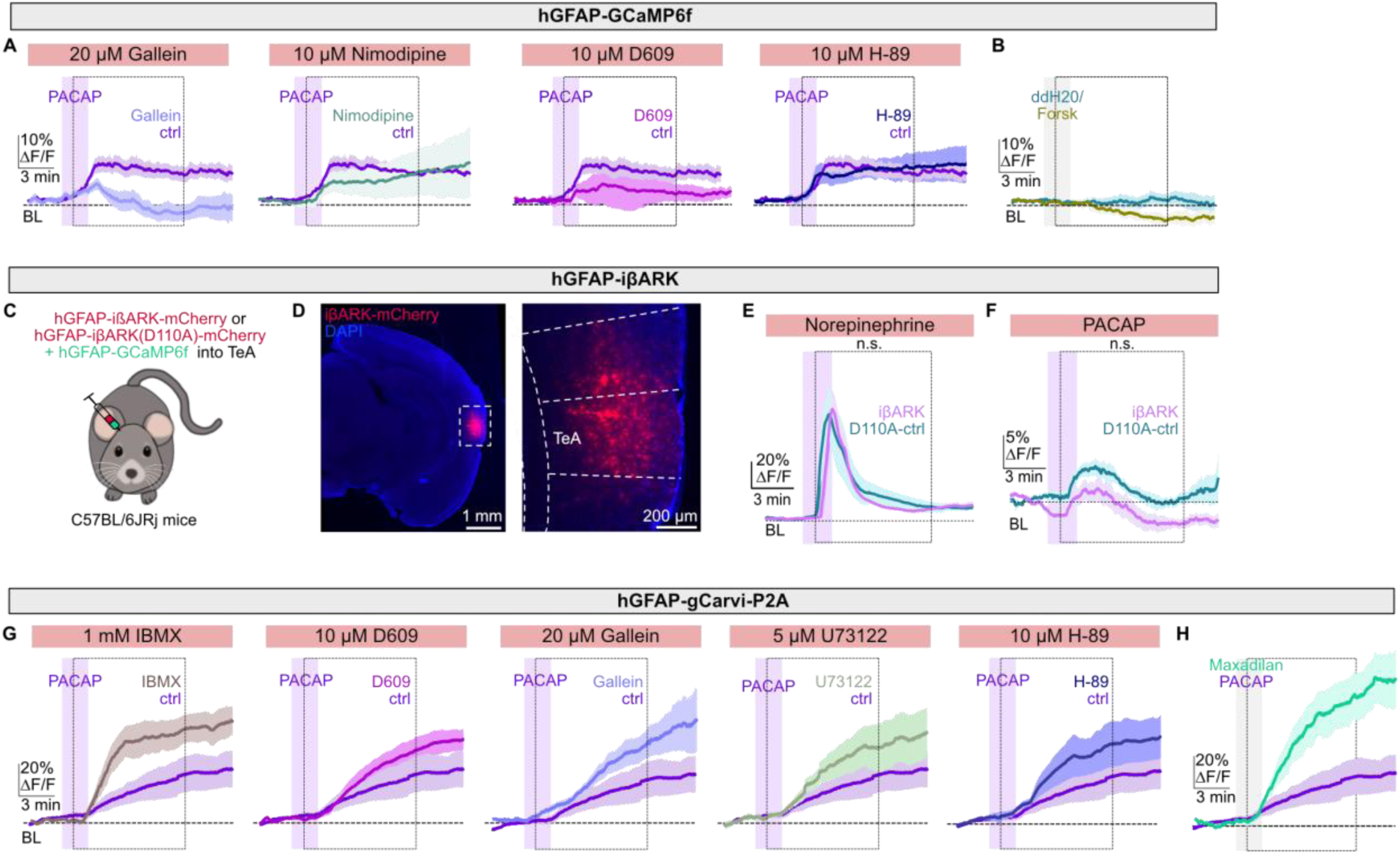
**A.** Averaged fluorescence Ca^2+^ signals (ΔF/F) from TeA astrocytes in response to PACAP application in the presence of 20 µM gallein, 10 µM nimodipine, 10 µM D609, or 10 µM H-89 compared to controls. Purple boxes indicate PACAP application. **B.** Averaged fluorescence Ca^2+^ signals (ΔF/F) from TeA astrocytes in response to 10 µM Forskolin application compared to ddH2O control (grey box). Forskolin: 2.07 ± 0.99% ΔF/F, *t*(10) = 0.68, *p* = 0.511; unpaired *t*-test, *N* = 6 slices per condition. **C.** Schematic of AAV-hGFAP-ißARK-mCherry and control virus with AAV-hGFAP-gCaMP6f injections into TeA. **D.** Exemplary epifluorescence image showing ißARK-mCherry expression. **E.** Averaged fluorescence Ca^2+^ signals (ΔF/F) in response to norepinephrine recorded from TeA astrocytes expressing either AAV-hGFAP-ißARK-mCherry (peak: 75.48 ± 12.11% ΔF/F) or AAV-hGFAP-ißARK-mCherry(D110A) (peak: 86.71 ± 20.49% ΔF/F), *t*(11) = 0.451, *p* = 0.661; unpaired *t*-test, *N* = 6-7 slices per condition. **F.** Averaged fluorescence Ca^2+^ signals (ΔF/F) in response to PACAP recorded from TeA astrocytes expressing either AAV-hGFAP-ißARK-mCherry (peak: 4.47 ± 1.72% ΔF/F) or AAV-hGFAP-ißARK-mCherry(D110A) (peak: 7.95 ± 1.14% ΔF/F), *t*(12) = 1.68, *p* = 0.119; unpaired *t*-test, *N* = 7 slices per condition. **G.** Averaged fluorescence cAMP signals (ΔF/F) from TeA astrocytes in response to PACAP application in the presence of 1 mM IBMX, 10 µM D609, 20 µM gallein, 5 µM U73122, or 10 µM H-89 compared to controls. Purple boxes indicate PACAP application. **H.** Averaged peak fluorescence cAMP signals (ΔF/F) from TeA astrocytes in response to PACAP and maxadilan (grey box). Peak signal PACAP: 21.19% [17.17-25.32%], maxadilan: 64.25 [46.58-75.32], *U* = 4.0, *p* = 0.026, Mann-Whitney *U* test, *N* = 6 slices per condition. Data are shown as mean ± SEM or median [IQR].

